# Forkhead box R1-mediated stress response linked to a case of human microcephaly and brain atrophy

**DOI:** 10.1101/2020.11.03.366740

**Authors:** Andressa Mota, Rui Hong, Sheng-Yong Niu, Féodora L. Bertherat, Lynne Wolfe, Christine May Malicdan, Thomas C. Markello, David R. Adams, William A. Gahl, Christine Cheng, Uwe Beffert, Angela Ho

## Abstract

Forkhead box (Fox) family transcription factors are highly conserved and play essential roles in a wide range of cellular and developmental processes. We report an individual with severe neurological symptoms including postnatal microcephaly, progressive brain atrophy and global developmental delay associated with a *de novo* missense variant (M280L) in the *FOXR1* gene. At the protein level, M280L impaired FOXR1 expression and induced a nuclear aggregate phenotype due to protein misfolding and proteolysis. RNAseq and pathway analysis showed that FOXR1 acts as both a transcriptional activator and repressor with central roles in heat shock response, chaperone cofactor-dependent protein refolding and cellular response to stress. Indeed, FOXR1 expression is increased in response to cellular stress, a process in which it directly controls *HSPA6, HSPA1A* and *DHRS2* transcripts. Meanwhile, the ability of the M280L mutant to respond to stress is compromised, in part due to impaired regulation of downstream target genes that are involved in the stress response pathway. Combined, these results suggest FOXR1 plays a role in cellular stress and is necessary for normal brain development.

## Introduction

Neurodevelopmental disorders result from abnormal brain development and the inability to reach cognitive, emotional and motor developmental milestones. Progress in genomics has advanced the prognosis of human neurodevelopmental disorders and provided insights into the molecular mechanisms of disease (McCarroll & Hyman, 2013; Tarlungeanu & Novarino, 2018; Parenti *et al*, 2020). While some causal genes are highly penetrant, there are also many rare single-nucleotide changes that have deleterious effects on genes of unknown function. Through exome sequencing, the NIH Undiagnosed Diseases Program (NIH UDP), a clinical site of the NIH Undiagnosed Diseases Network (UDN) identified a variant (M280L) in a single allele of the *FOXR1* gene (forkhead box R1; NM_181721.2) in an individual with severe neurological symptoms including postnatal microcephaly, progressive brain atrophy and global developmental delay.

FOXR1 is a member of the evolutionarily conserved forkhead box (Fox) family of transcription factors named after the ectopic head structures observed in mutants of the *Drosophila* gene *forkhead* (*fkh*) (Weigel *et al*, 1989; Kaestner *et al*, 2000; Mazet *et al*, 2003). Mutations in the *Drosophila fkh* gene cause defects in head fold involution during embryogenesis, resulting in a characteristic spiked head appearance in adult flies. Since the discovery of *fkh*, hundreds of *Fox* genes have been identified in organisms ranging from yeasts to humans, making it one of the largest but least explored families of higher eukaryotic transcription factors (reviewed in Hannenhalli & Kaestner, 2009; Golson & Kaestner, 2016). All members of the *Fox* gene family of transcription factors are monomeric, helix-turn-helix proteins that harbor a core fkh DNA-binding domain comprised of three α-helices connected *via* a small β-sheet to a pair of loops resembling butterfly wings or a “winged-helix” (Clark *et al*, 1993; Gajiwala & Burley, 2000; van Dongen *et al*, 2000). Despite the high degree of conservation identity in the DNA-binding domain, Fox proteins bind different target sequences with great specificity, effecting regulation of the transcription of large array of genes directing major developmental processes such as cell proliferation and cell fate specification (Clark *et al*., 1993; Carlsson & Mahlapuu, 2002; Lehmann *et al*, 2003; Nakagawa *et al*, 2013). Human genetic analyses have shown several *FOX* genes have important biological functions associated with brain development; these include *FOXG1* (potential determinant of forebrain size; Florian *et al*, 2012; Hettige & Ernst, 2019; Pringsheim *et al*, 2019) and *FOXP2* (vocal learning; MacDermot *et al*, 2005; Fisher & Scharff, 2009; Nudel & Newbury, 2013). Further, mutations in *FOXG1, FOXC2, FOXL2, FOXP1* and *FOXP2* have profound effects on human brain development including microcephaly, intellectual impairments, and language disorders (D’Haene *et al*, 2010; Kortum *et al*, 2011; Butler *et al*, 2012; Seltzer & Paciorkowski, 2014; Han *et al*, 2019).

FOXR1, also known as FOXN5 (forkhead box N5) or DLNB13, is a 292 amino acid protein that contains an fkh DNA-binding domain (Katoh and Katoh, 2004a). Human *FOXR1* and rat *foxr1* gene consist of six exons with conserved exon-intron structure, indicating that FOXR1 is well-conserved between human and rat genomes (Katoh and Katoh, 2004b). The Genome-based tissue expression consortium indicate that, *FOXR1* is expressed in the human brain and reproductive organs (Consortium, 2013). The Human Brain Transcriptome shows that *FOXR1* is expressed in all brain regions during embryonic and postnatal development and its expression level in the brain is maintained throughout life (https://hbatlas.org). Furthermore, *in situ* hybridization showed that mouse *foxr1* expression was present in all brain regions and enhanced within cellular nuclei, consistent with the human tissue expression profile based on the Allen Brain Atlas (Lein *et al*, 2007). However, little is known about the function of FOXR1. Several studies have shown that mouse *foxr1* is involved in spermiogenesis (Petit *et al*, 2015), and several point mutations within human *FOXR1* have been shown to be associated with a variety of carcinomas, although functional characterization of these oncogenic *FOXR1* mutants have not been performed (Santo *et al*, 2012; Katoh, 2013; Pommerenke *et al*, 2016). Recently, *foxr1* was found to be an essential maternal– effect gene in zebrafish and is required for proper cell division and survival (Cheung *et al*, 2018).

Here, we report a human neurodevelopmental disorder associated with a rare variant in *FOXR1*. We demonstrate that the *de novo* missense M280L variant decreases FOXR1 protein expression and displayed nuclear puncta aggregates in HEK293T cells, suggesting that impaired FOXR1 function can be pathogenic. In addition, we provide evidence that the FOXR1 M280L mutant has a compromised ability to respond to stress, in part due to impaired regulation of downstream target genes that are involved in the stress response pathway.

## Results

### Exome sequencing identified an individual with global developmental delay carrying a *de novo* missense variant in *FOXR1*

The NIH UDP identified a proband with severe neurological symptoms including postnatal microcephaly, progressive brain atrophy and severe muscle hypotonia from early infancy. Brain MRI showed progressive hypoplasia in the cerebral cortex, pons and cerebellum and ventricular enlargement from age 1 to 5 (Fig 1A). The proband also exhibited growth delay, decreased body weight, short stature, scoliosis, hip dysplasia, ankle clonus and bell-shaped thorax (Appendix Table 1). Ophthalmic abnormalities included particular optic atrophy, cortical visual impairment and retinitis pigmentosa. Neuromuscular abnormalities included hyperactive deep tendon reflexes, joint hypermobility, severe muscle hypotonia and poor head control. The proband had myopathic facies, preauricular pits, anteverted nares and low set ears.

**Figure 1.**
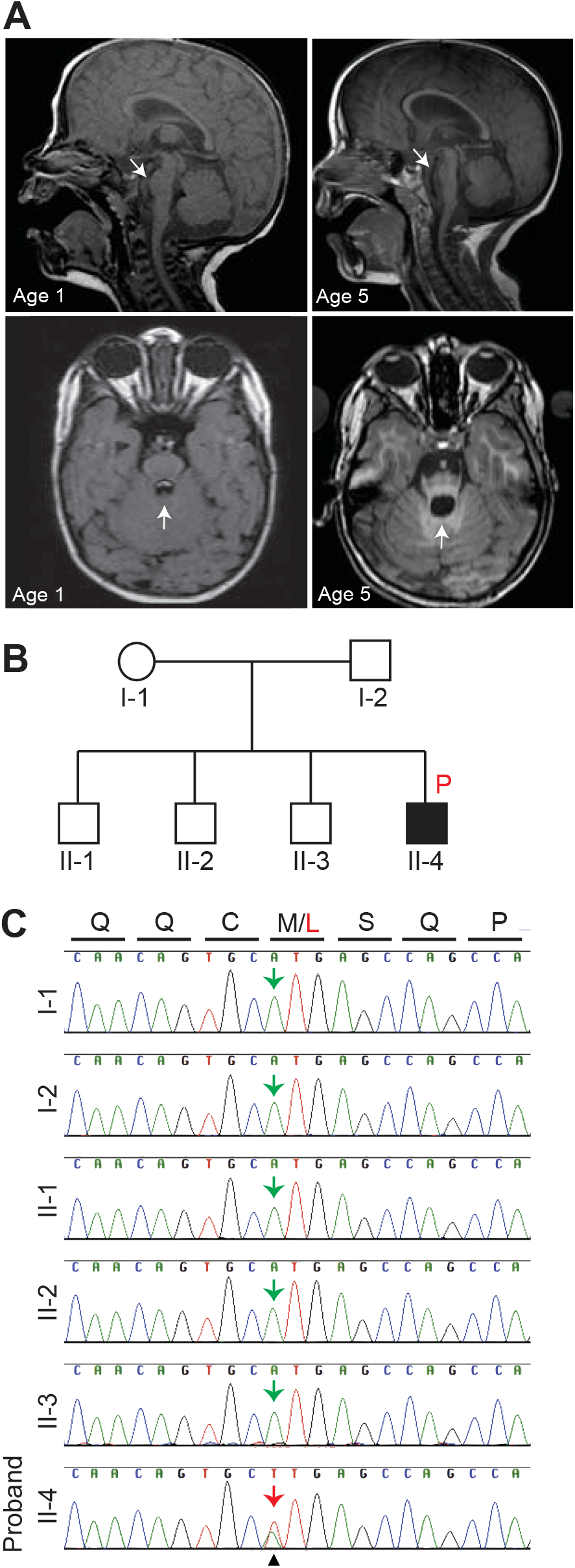
Neuroimaging and identification of a *de novo FOXR1* missense variant in a proband with microcephaly and brain atrophy. A. Top images, MRI of mid-sagittal images of the proband at 1 and 5 years old showing hypoplasia of the pons (arrows) and cerebellum. Bottom images, Horizontal view of the proband at 1 and 5 years old showing dilation of ventricle indicated by the arrows. B. Pedigree of the family where the letter P in red (black square) indicates the proband. C. Sanger sequence analysis confirming the *de novo FOXR1* variant. Sequence chromatograms demonstrate the presence of the heterozygous variant in the proband, II-4 (indicated by the red arrow) and the reference allele in both parents and siblings (green arrows). Letters on top indicate amino acid residues (Q = glutamine, C = cysteine, M = methionine, L = leucine, S = serine, P = proline).

Exome sequencing was performed on the proband and the siblings and parents who are all unaffected. Three likely pathogenic candidate genes, rapamycin and FKBP12 target (*RAFT1*), ATPase Na^+^/K^+^ transporting subunit alpha 3 (*ATP1A3*), and *FOXR1* were identified. RAFT1 functions as a kinase that regulates cell growth, proliferation, motility, and survival (Sabatini *et al*, 1994; Burnett *et al*, 1998). The proband had a homozygous *RAFT1* missense mutation, but the EXAC database identified an unaffected individual with the same *RAFT1* mutation. The second candidate, *ATP1A3*, maintains plasma membrane sodium and potassium gradients (Heinzen *et al*, 2014). Investigations revealed an individual with the same mutation was discovered and a mild phenotype involving learning disability and episodes of dizziness. The last candidate is a *de novo* missense variant in *FOXR1*, a gene of unknown function, and the variant was not identified in the siblings and parents (Fig 1B). The heterozygous *de novo* nonsynonymous variant results in a methionine-to-leucine substitution at position 280 (M280L) and was confirmed by Sanger sequencing (Fig 1C). The M280 is found in the C-terminal segment of the FOXR1 protein, which is outside of the DNA-binding domain. M280 is highly conserved through evolution, from mammals, birds, reptiles to frogs and zebrafish (Appendix Fig 1). In addition, the M280L variant is predicted to be damaging and disease-causing based on scores of Combined Annotation Dependent Depletion (score of 29.9 where a score of 30 means that the variant is in the top 0.1% of deleterious variants in the human genome), PolyPhen-2 (score: 0.994/1.0), and Mutation Taster (score: 0.99/1.0).

### The FOXR1 M280L mutant leads to a decrease in FOXR1 protein expression

To examine whether the FOXR1 M280L mutant was properly expressed *in vitro*, we transiently transfected FOXR1 wild-type (WT) or the M280L mutant in HEK293T or COS7 cells and immunoblotted for FOXR1 or GFP-tagged FOXR1 protein. FOXR1 levels were significantly decreased by the M280L mutant (Fig 2A-C). Since *FOXR1* is a transcription factor, we next tested whether the M280L mutant affected FOXR1 nuclear distribution in HEK293T cells transfected with either untagged or GFP-tagged FOXR1 WT or M280L. Western blot analysis demonstrated that both FOXR1 WT and M280L protein was expressed in both cytoplasmic and nuclear fractions with higher expression found in the nuclear fraction (Fig 2D). However, expression of the M280L mutant was reduced compared to FOXR1 WT in both cytoplasmic and nuclear fractions.

**Figure 2.**
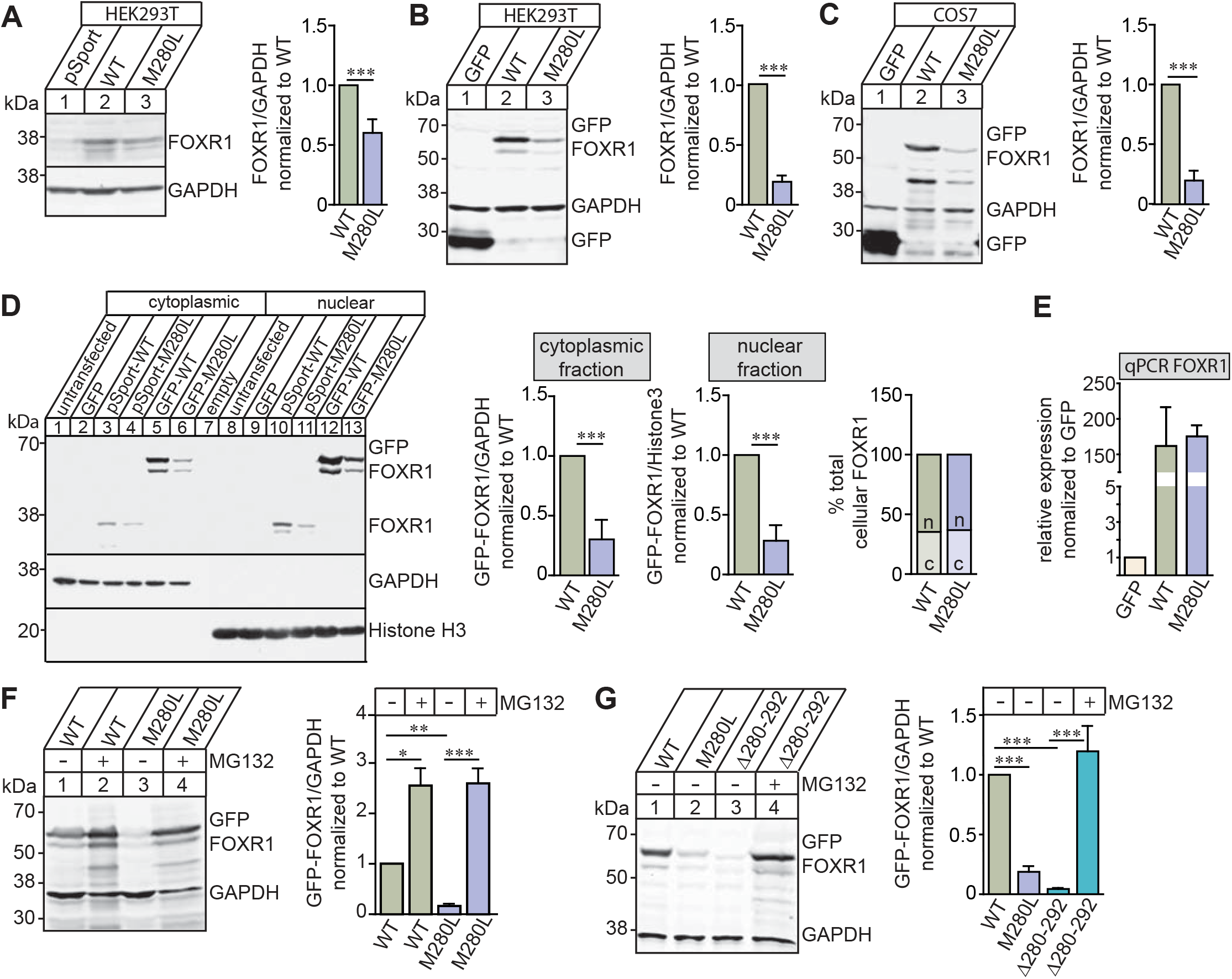
The M280L mutant destabilizes FOXR1 protein. A. Representative immunoblots and quantitative analysis of FOXR1 from HEK293T cells transfected with pCMV-SPORT6 human FOXR1 WT or M280L mutant. GAPDH served as a loading control. Graph represents FOXR1 over GAPDH normalized to WT. Unpaired *t*-test (n = 6 independent experiments, *** p < 0.0001). B. Representative immunoblot and quantitative analysis of FOXR1 from HEK293T cells transfected with GFP, GFP-tagged human FOXR1 WT or M280L mutant. GAPDH served as a loading control. Graph represents FOXR1 over GAPDH normalized to WT. Unpaired *t*-test (n = 5 independent experiments, *** p < 0.0001). C. Representative immunoblot and quantitative analysis of FOXR1 from COS7 cells transfected with GFP, GFP-tagged human FOXR1 WT or M280L mutant. GAPDH served as a loading control. Graph represents FOXR1 over GAPDH normalized to WT. Unpaired *t*-test (n = 4 independent experiments, ** p = 0.0013). D. Representative immunoblots and quantitative analysis of cytoplasmic (c) and nuclear (n) fractions of FOXR1 from HEK293T cells transfected with pCMV-SPORT6 or GFP-tagged human FOXR1 WT and M280L. GAPDH and Histone H3 served as cytoplasmic and nuclear loading markers, respectively. Graph represents FOXR1 over GAPDH normalized to WT. Unpaired *t*-test (n = 5 independent experiments, *** p < 0.0001). The percentages of total cellular FOXR1 is the cytoplasmic and nuclear fractions were determined. E. Quantitative PCR (qPCR) to quantify *FOXR1* mRNA levels from HEK293T cells transfected with GFP, GFP-tagged human FOXR1 WT or M280L mutant. Graph represents relative *FOXR1* mRNA expression normalized to GFP. F. Representative immunoblot and quantitative analysis of FOXR1 from HEK293T cells transfected with GFP-tagged human FOXR1 WT or M280L mutant. Protein stability was monitored by quantitative immunoblotting after blocking with proteasome inhibitor MG132. Graph represents FOXR1 over GAPDH normalized to untreated WT. One-way ANOVA (n = 4 independent experiments, * p = 0.0143, ** p = 0.0012, *** p = 0.0003). G. Representative immunoblot and quantitative analysis of FOXR1 from HEK293T cells transfected with GFP-tagged human FOXR1 WT, M280L mutant or FOXR1 C-terminal truncation mutant lacking the last 12 amino acids (Δ280-292). Protein stability was monitored for FOXR1 Δ280-292 mutant by blocking with MG132. GAPDH served as a loading control. Graph represents FOXR1 over GAPDH normalized to untreated WT. One-way ANOVA (n = 3 independent experiments, *** p < 0.0001).

We next investigated whether the decrease in FOXR1 expression in the M280L mutant was due to transcription or protein stability changes. In HEK293T-transfected cells, we detected equal amounts of *FOXR1* mRNA levels of FOXR1 WT and M280L, indicating that decreased M280L protein expression was not due to decreased transcription (Fig 2E). To measure protein stability, we blocked the proteasome pathway by treating transfected HEK293T cells with MG132, a cell-permeable proteasome inhibitor. Protein levels of both FOXR1 WT and M280L were approximately the same after proteasome inhibition, suggesting that the M280L mutant destabilizes FOXR1 protein, likely due to protein misfolding making it susceptible to proteolysis and degradation through the proteasome pathway (Fig 2F).

Finally, we investigated whether the short C-terminal tail containing M280 is necessary for protein stabilization. We produced a FOXR1 C-terminal truncation mutant lacking the last 12 amino acids from M280 (Δ280-292). Indeed, HEK293T cells transfected with GFP-tagged Δ280-292 had decreased FOXR1 protein levels, which increased following MG132 treatment, suggesting that the FOXR1 C-terminal tail is critical for FOXR1 protein stability (Fig 2G).

### FOXR1 M280L induces a nuclear aggregate phenotype

To examine whether the M280L mutant alters the cellular localization of FOXR1, we transfected HEK293T cells with GFP, GFP-tagged FOXR1 WT, M280L or Δ280-292 mutant. Immunostaining for GFP showed expression of FOXR1 WT mainly in a diffuse pattern in the nucleus, co-localizing with DAPI nuclear marker (Fig 3A). In contrast, about 13% of cells transfected with the M280L mutant formed discrete nuclear puncta (Fig 3B-C). We observed a similar phenotype in COS7 cells transfected with the M280L variant (Appendix Fig 2). In nuclei containing >15 puncta, the average size of individual puncta was <2 μm^2^, whereas nuclei containing <5 puncta had aggregates of >4 μm^2^ (Fig 3D). These results suggest that the larger puncta may form by coalescing from small nuclear foci. Expression of FOXR1 Δ280-292 mutant displayed a similar pattern, suggesting that the nuclear puncta could be due to misfolded FOXR1 protein that is susceptible to aggregation.

**Figure 3.**
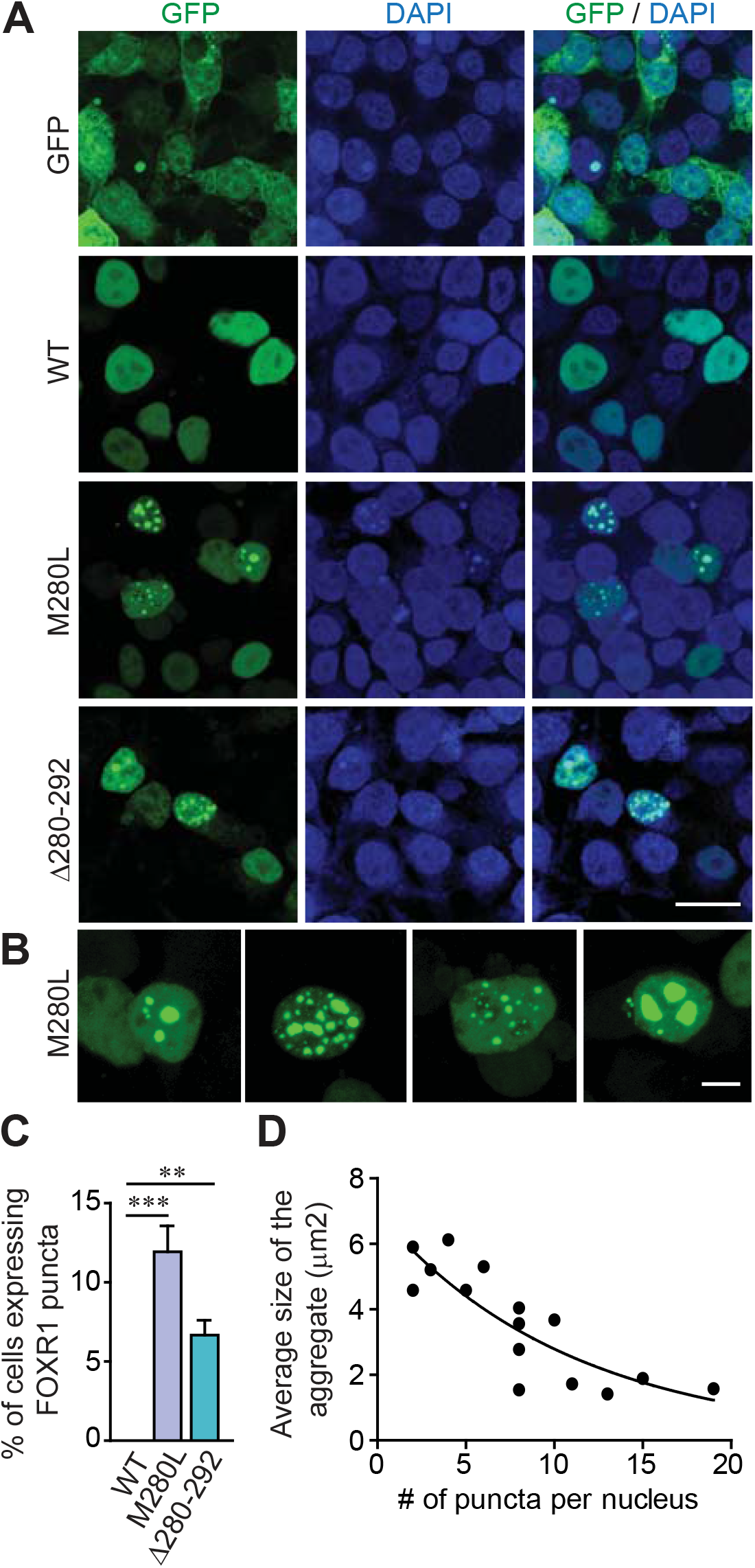
The M280L variant induced nuclear puncta phenotype. A. Fluorescence images of HEK293T cells transfected with GFP or GFP-tagged human FOXR1 WT, M280L or Δ280-292 mutants. DAPI (blue) served as a nuclear marker. Scale bar = 20 μm. B. Fluorescence images of HEK293T cells transfected with GFP-tagged M280L mutant showing a range of nuclear puncta phenotypes. Scale bar = 5 μm. C. Quantitative analysis on the percentage of cells expressing FOXR1 puncta phenotype. One-way ANOVA (n = 3 independent experiments, **p = 0.0048, *** p = 0.0002). D. Correlation analysis of the average size of the aggregate to the number of puncta per nucleus.

### Identification of novel FOXR1-dependent transcripts by RNA sequencing analysis

To identify target genes regulated by FOXR1 and to investigate the role of FOXR1 M280L mutant, we performed an unbiased transcriptomic screen by RNA sequencing (RNAseq) in HEK293T cells transiently transfected with GFP, GFP-tagged FOXR1 WT or M280L. Principal component analysis showed that the three groups clustered separately excluding experimental covariates and batch effects (Appendix Fig 3A). We plotted a heat map of the log (2) fold change for all the differentially-expressed genes (DEGs) and delineated five coherent clusters (Fig 4A). Differential gene expression analysis of FOXR1 WT transfected cells compared to GFP identified 2644 DEGs of which 1315 (49.7%) were upregulated and 1329 (50.3%) were downregulated transcripts, suggesting that *FOXR1* acts as both a transcriptional activator and repressor (Fig 4A, Appendix Fig 3B). When we compared WT to M280L, we identified 735 DEGs of which 561 (76.3%) were upregulated and 174 (23.7%) genes were downregulated (Appendix Fig 3B). We paid special attention to those transcripts whose levels showed a 2-fold increase in FOXR1 WT and a decrease in M280L relative to GFP control as delineated in cluster E (Fig 4B).

**Figure 4.**
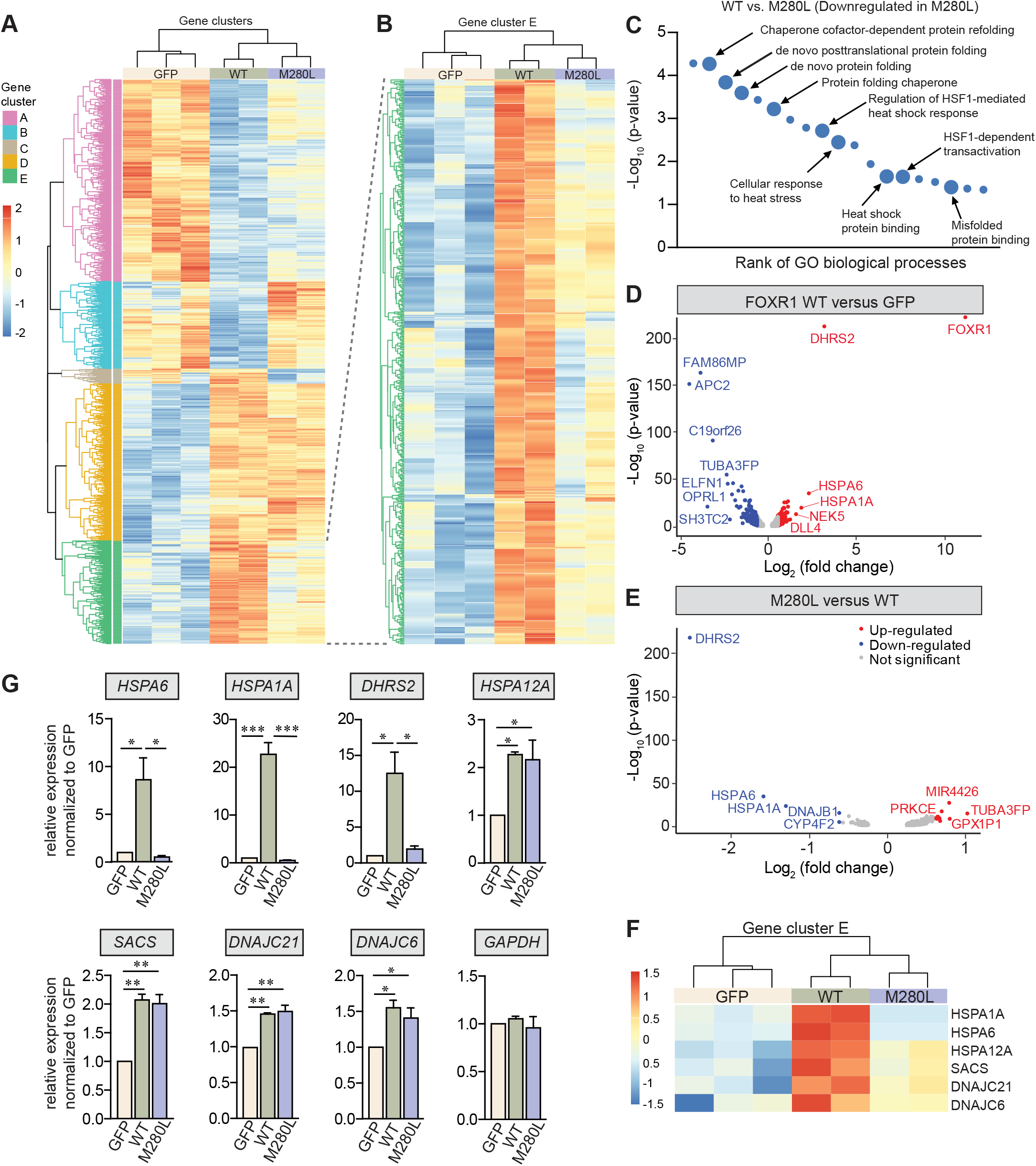
RNAseq analysis of FOXR1 wild-type and M280L mutant. A. Heatmap of hierarchical clustering indicates differentially-expressed genes (rows) between GFP, GFP-tagged FOXR1 WT and M280L (fold-change > 2, p < 0.05). Red indicates up-regulated genes and blue indicates down-regulated genes. B. Heatmap of gene cluster ‘E’ indicates differentially-expressed genes (rows) that are upregulated in FOXR1 WT and down-regulated in M280L. C. Distribution of gene ontology (GO) terms annotated in biological processes of highly-regulated genes in FOXR1 WT and down-regulated in M280L. D. Volcano plot of differentially expressed genes that were up-regulated between FOXR1 WT versus GFP control. Significantly up-regulated genes are in red while down-regulated genes are in blue. Non-significant genes are in gray. E. Volcano plot of differentially expressed genes that were down-regulated between FOXR1 M280L versus WT. Significantly down-regulated genes are in blue and up-regulated are in red. Non-significant genes are in gray. F. Heatmap of gene cluster ‘E’ highlighting several chaperone proteins that were differentially expressed in FOXR1 WT and down-regulated in M280L. G. Quantitative real-time PCR verifying the RNAseq analysis showing FOXR1 drives expression of *HSPA6, HSPA1A* and *DHRS2* and are misregulated in the M280L mutant. Graph represents relative expression normalized to GFP. One-way ANOVA (n = 3 independent experiments, * p < 0.05, ** p<0.005, *** p < 0.0001).

Gene ontology (GO) analysis for biological processes within cluster E showed genes involved in the heat shock response. These were functionally-related to negative regulation of inclusion body assembly, chaperone cofactor-dependent protein refolding, *de novo* protein folding, cellular response to stress and regulation of HSF1-mediated heat shock response where these were enriched in FOXR1 WT and downregulated in M280L (Fig 4C and Appendix Fig 4). Based upon the volcano plots that summarize both the expression fold-change and the statistical significance, the upregulated genes in response to FOXR1 WT and downregulated in M280L included *HSPA1A* and *HSPA6* (both members of the Hsp70 family of heat shock proteins, Hsps), and *DHRS2* (Dehydrogenase/Reductase SDR Family Member 2, a mitochondrial reductase enzyme) (Fig 4D-E). These proteins play roles in protecting against oxidative stress-mediated cellular response. Quantitative real-time-PCR (qRT-PCR) supported the RNAseq data for *HSPA6, HSAPA1A* and *DHRS2* (Fig 4F-G), confirming upregulation of gene expression in FOXR1 WT and not in the M280L mutant. Other Hsps such as *SACS, DNAJC21* and *DNAJC6* were increased in both FOXR1 WT and M280L groups (Fig 4G). Not all members of the Hsp70 family were misregulated in the M280L mutant; for example, *HSPA12A* transcript was found to be upregulated in both FOXR1 WT and the M280L mutant (Fig 4G). These results indicate that FOXR1 drives expression of specific Hsps and an important NADPH-dependent reductase enzyme that is likely related to cytoprotective pathways alleviating oxidative stress.

### FOXR1 controls gene expression of heat shock chaperones and an anti-oxidant NADPH-dependent reductase

To determine whether *HSPA6, HSPA1A* and *DHRS2* are directly regulated by FOXR1 as opposed to secondary targets of FOXR1, we performed a *de novo* motif analysis of target promoters to identify consensus DNA-binding sites (Appendix Fig 5). We found strong consensus sequences for FOXR1 response elements (Nakagawa *et al*., 2013) within the promoter regions of at least three of the top FOXR1-regulated genes, *HSPA6, HSPA1A* and *DHRS2* (Fig 5A). To determine whether FOXR1 expression regulates the activity of these promoters, a dual luciferase system under the control of proximal upstream regions of human *HSPA6* (−1119 to -113 bp), *HSPA1A* (−1053 to -210 bp) and *DHRS2* (−3329 to -2313 bp) were co-transfected with either GFP control, FOXR1 WT or M280L mutant in HEK293T cells (Fig 5B). We found that *HSPA6, HSPA1A* and *DHRS2* were activated by FOXR1 WT but not by M280L, indicating that these promoter regions contain FOXR1 responsive sequences and are direct targets of FOXR1 WT. The lack of luciferase activity observed in the M280L mutant is likely due to the decrease in FOXR1 expression levels (Fig 5C).

**Figure 5.**
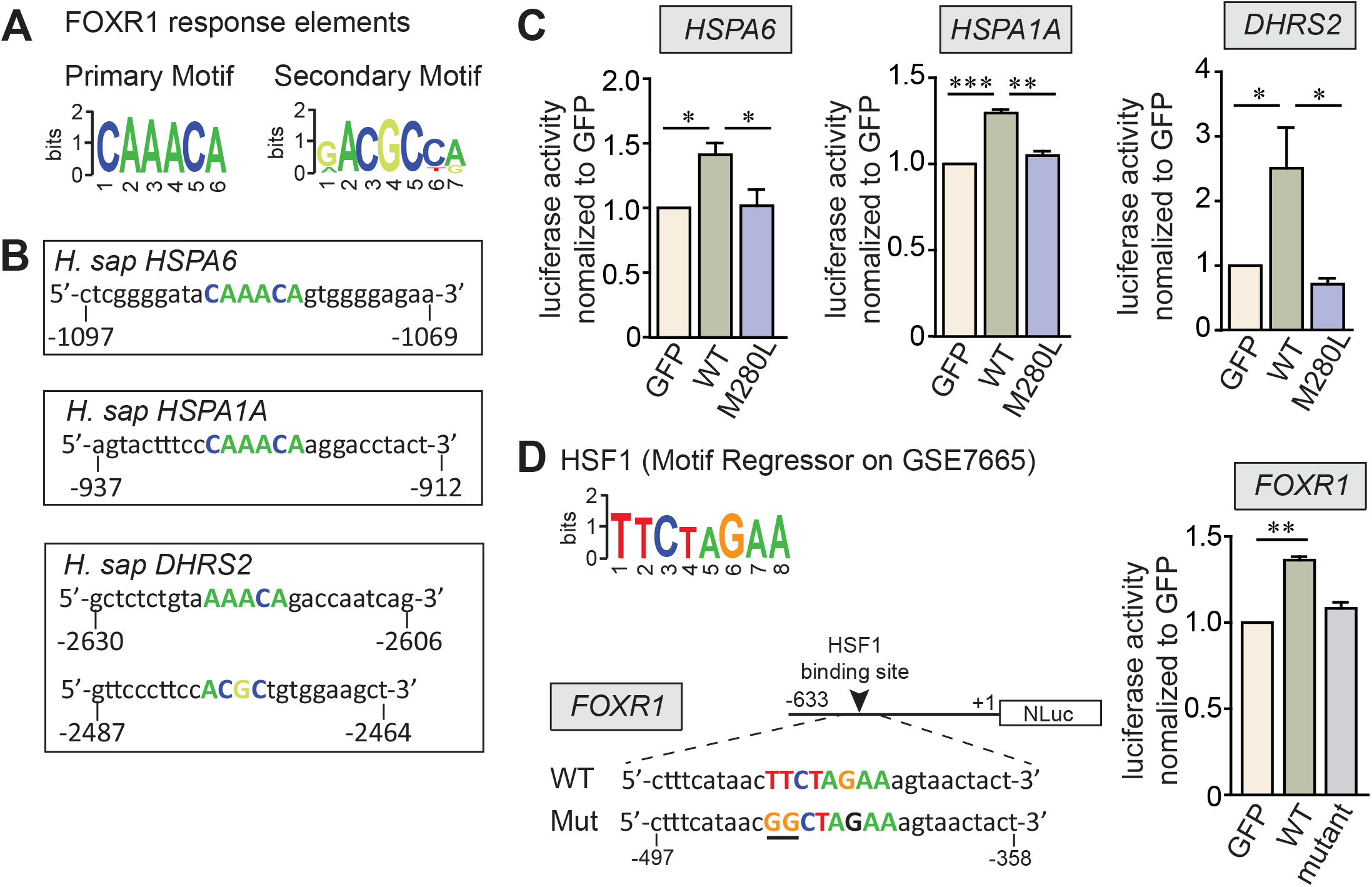
Human DNA binding-site motifs bound by FOXR1. A. FOXR1 response elements showing consensus primary and secondary sequences bound by FOXR1 (adapted from Nakagawa *et al*., 2013). B. Putative FOXR1 response elements are denoted in the promoters of three of the top-regulated FOXR1-targeted genes: *HSPA6, HSPA1A* and *DHRS2*. C. Dual luciferase reporter assays where GFP control, FOXR1 WT or M280L were co-transfected into HEK293T cells with the corresponding *HSPA6, HSPA1A* and *DHRS2* luciferase reporters. Data are plotted as luciferase activity normalized to GFP control. One-way ANOVA (n = 3 independent experiments, * p < 0.05, ** p<0.005, *** p < 0.0001). D. Consensus primary sequences bound by HSF1. The putative HSF1 response elements are denoted in the promoter of *FOXR1*. Dual luciferase reporter assays in HEK293T cells comparing *FOXR1* promoter activation to a *FOXR1* mutant where two residues of the HSF1 response elements in FOXR1 were mutated from TT to GG (mutant). Data was plotted as luciferase activity normalized to GFP control. One-way ANOVA (n = 3 independent experiments, ** p = 0.0062).

Expression of Hsps is known to be regulated by the transcription factor heat shock factor 1 (HSF1), which has a high affinity for *cis*-acting DNA sequence elements, including the heat shock elements (HSEs) found in the promoters of HSF-responsive genes such as Hsp70 proteins (reviewed in Akerfelt *et al*, 2010). There is also precedence that HSF1 target genes extend beyond molecular chaperones. For example, in *C. elegans*, the protective effects of reduced insulin signaling require both HSF1 and the FOXO transcription factor, *DAF-16*, to prevent damage by protein misfolding and to promote longevity (Hsu *et al*, 2003; Morley & Morimoto, 2004; Singh & Aballay, 2009). Based on the GO analysis for biological processes, transcripts that were upregulated in FOXR1-transfected cells were genes related to regulation of HSF1-mediated heat shock response (Appendix Fig 4). Hence, we searched and identified FOXR1 response elements upstream of the HSEs in both the *HSPA6* and *HSPA1A* promoters (Appendix Fig 5). We also identified a consensus sequence for binding by HSF1 within the promoter region of FOXR1. Utilizing a dual luciferase system under the control of an upstream region of human FOXR1 (−633 to +1 bp), FOXR1 was found to be activated by HSF1. However, HSF1-mediated FOXR1 activation was not observed when the HSF response element was mutated from TTCTAGAA to GGCTAGAA *in vitro*, indicating that human FOXR1 is a direct target of HSF1, which may be regulated by cellular stress (Fig 5D).

### FOXR1 expression is increased in response to cellular stress

Because FOXR1 regulates expression of *HSPA6* and *HSPA1A* transcripts and they are also direct targets of HSF1, we hypothesized that FOXR1 expression might be directly regulated following stress-induced paradigms. We induced cellular stress using two different paradigms: serum deprivation (metabolic stress for 24 hours) and CO_2_-deprivation (oxidative stress for 24 hours). Under both paradigms, cells transfected with FOXR1 WT exhibited a 2.5- and 3.3-fold increase in FOXR1 protein expression under serum- and CO_2_-deprivation, respectively when compared to the non-stressed condition (Fig 6A). The increase in FOXR1 protein levels coincided with an increase in nuclear FOXR1 (Fig 6B). In contrast, FOXR1 M280L expression also exhibited a 3.3-fold increase under CO_2_-deprivation but not during serum-deprivation, indicating that the M280L mutant may be sensitive to different types of environmental stressors. In fact, the number of nuclear aggregates in cells transfected with the M280L mutant in response to CO_2_-deprivation was increase but not in response to serum-deprivation (Appendix movie 1).

**Figure 6.**
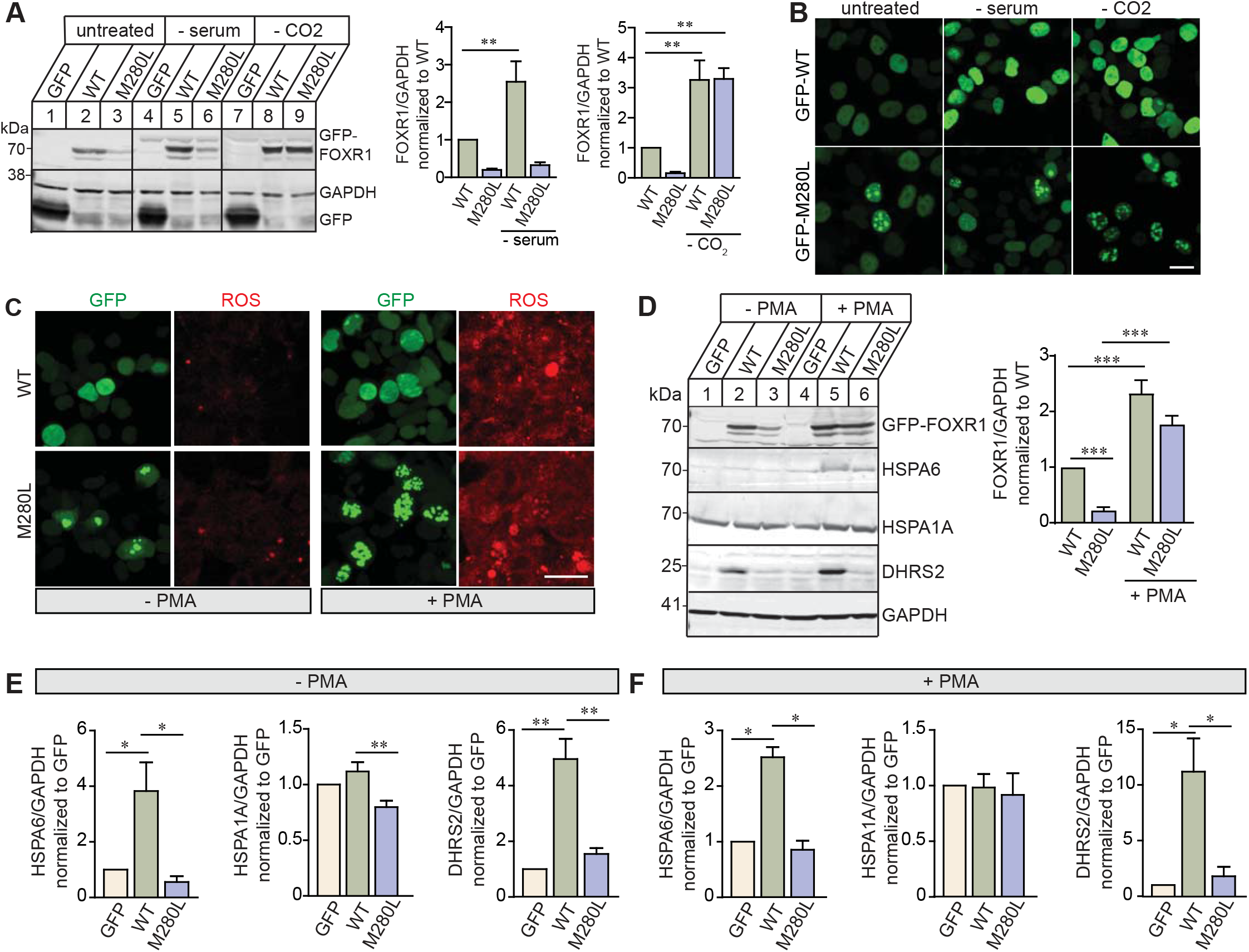
FOXR1 expression is increased in response to cellular stress. A. Representative immunoblots and quantitative analysis for FOXR1 from HEK293T cells transfected with GFP, GFP-tagged FOXR1 WT or M280L mutant in response to serum and CO_2_ deprivation. GAPDH served as loading control. Graph represents FOXR1 over GAPDH normalized to untreated WT. One-way ANOVA (n = 4 independent experiments, ** p <0.005). B. Fluorescence images of HEK293T cells transfected with GFP-tagged human FOXR1 WT or M280L in response to serum and CO_2_ deprivation. Scale bar = 20 μm. C. Fluorescence images of HEK293T cells transfected with GFP-tagged human FOXR1 WT or M280L and treated with PMA, a NADPH oxidase activator known to enhance reactive oxygen species (ROS). Cells were fixed after 24 hours of treatment and assessed for ROS generation using CellROX, a photostable ROS sensor. Scale bar = 20 μm. D. Representative immunoblots and quantitative analysis of HEK293T cells following PMA treatment showing an increase in FOXR1 expression. Graph represents FOXR1 over GAPDH normalized to untreated WT. One-way ANOVA (n = 5 independent experiments, *** p<0.0005). E. Quantitative analysis of HSPA6, HSPA1A and DHRS2 protein levels from HEK293T cells transfected with GFP, GFP-tagged human FOXR1 WT or M280L. Graph represents protein of interest over GAPDH normalized to GFP. One-way ANOVA (n = 4-6 independent experiments, * p < 0.05, ** p<0.005). F. Quantitative analysis of HSPA6, HSPA1A and DHRS2 protein levels from HEK293T cells transfected with GFP, GFP-tagged human FOXR1 WT or M280L and treated with PMA. Graph represents protein of interest over GAPDH normalized to GFP. One-way ANOVA (n = 2-3 independent experiments, * p < 0.05).

To further explore the relationship between FOXR1 and oxidative stress, we treated FOXR1-transfected HEK293T cells with phorbol 12-myristate 13-acetate (PMA), a pharmacologic NADPH oxidase activator known to enhance reactive oxygen species (ROS) generation through a protein kinase C-mediated pathway (Bhat *et al*, 2019). We assessed ROS generation by fluorescence imaging using CellROX, a photostable ROS sensor. Consistent with other stress paradigms, PMA enhanced ROS generation in HEK293T cells transfected with FOXR1 WT and M280L (Figure 6C). PMA enhanced the diffuse FOXR1 expression in the nucleus of HEK293T cells transfected with FOXR1 WT. Also, the number of nuclear aggregates in cells transfected with the M280L mutant was increase by 3.88-fold compared to non-PMA treatment (two-tailed, *t* = 6.382 df = 9, p<0.0001), suggesting ROS-induced aggregation of FOXR1 protein in response to stress (Appendix movie 2). To determine whether ROS-induced aggregation of FOXR1 protein is cytotoxic, we measured the amount of lactate dehydrogenase (LDH) released into the medium as a measure of cytotoxicity and found no LDH changes across cells transfected with GFP alone and GFP-tagged FOXR1 WT (one-way ANOVA, F_(3,8)_=0.2963, p=0.6944) or FOXR1 WT compared to M280L (one-way ANOVA, F_(3,8)_=0.2963, p=0.8272), indicating that the nuclear aggregates were not cytotoxic.

We found FOXR1 protein expression was increased 2.3- and 1.8-fold in cells transfected with FOXR1 WT and M280L mutant after PMA treatment, respectively (Fig 6D). Concomitantly, we found an increase in both HSPA6 and DHRS2 protein expression levels in cells transfected with FOXR1 WT (Fig 6E-F). HSPA6 levels were increased in response to PMA treatment in the M280L mutant. In contrast, we did not observe any changes in DHRS2 protein expression levels in cells transfected with M280L regardless of PMA treatment. In addition, while we observed a significant increase in *HSPA1A* mRNA levels in cells transfected with FOXR1 WT (Fig 4), we did not detect any changes in HSPA1A protein levels in cells transfected with FOXR1 WT or M280L. However, we did consistently see a decrease in HSPA1A protein levels in cells transfected with M280L compared to FOXR1 WT, but this difference disappeared when cells were treated with PMA.

### FOXR1 nuclear puncta in M280L mutant are insoluble

To determine whether the nuclear puncta that form in HEK293T cells transfected with M280L mutant were aggresomes, which are known to serve as storage bins of misfolded or aggregated proteins (Shen *et al*, 2011), transfected HEK293T cells were treated with PMA and stained with the Proteostat dye. The dye detects misfolded and aggregated proteins in cells. We found bright punctate staining for proteostat-positive aggregates colocalized with the nuclear puncta in cells expressing the M280L mutant but not in FOXR1 WT (Fig 7A). These results were similar in transfected cells expressing M280L that were treated with the cell-permeable proteasome inhibitor MG132, further supporting that the M280L mutation destabilized FOXR1 protein and formed nuclear aggregates (Fig 7B).

**Figure 7.**
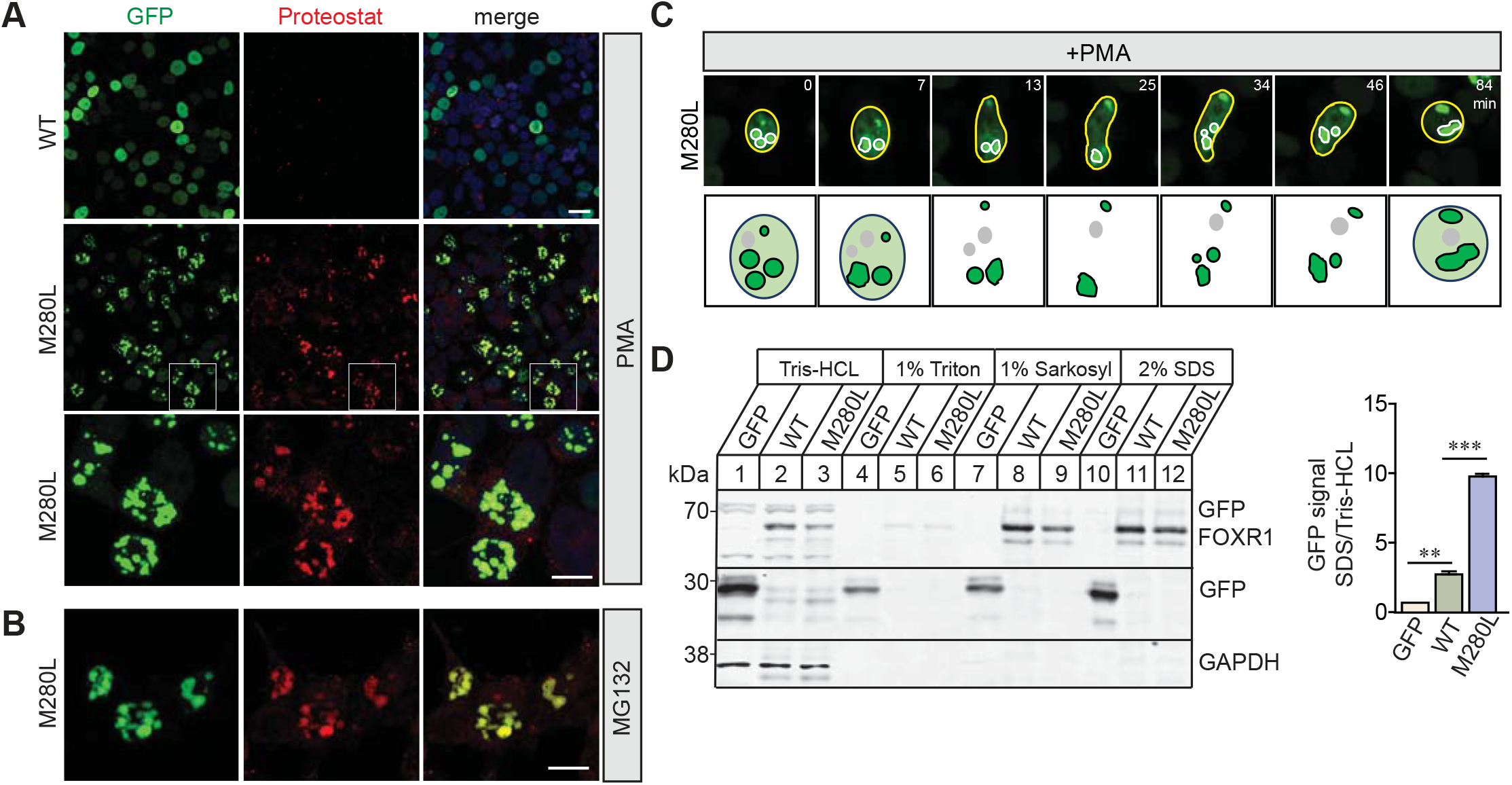
M280L nuclear aggregates are insoluble misfolded proteins. A. Fluorescence images of HEK293T cells transfected with GFP-tagged human FOXR1 WT or M280L and treated with PMA. Cells were fixed after 24 hours of treatment and immunolabel with Proteostat marker. White square box in the middle panels indicate images presented in the bottom panel at higher magnification. Top and middle panels, scale bar = 20 μm. Bottom panels, scale bar = 10 μm. B. Fluorescence images of HEK293T cells transfected with GFP-tagged M280L and treated with MG132. Cells were fixed after 24 hours of treatment and immunolabel with Proteostat marker. Scale bar = 10 μm. C. Time-lapse imaging of HEK293T cells transfected with GFP-tagged M280L. Top panel represent images showing nuclear aggregates undergoing extensive movements and fusions. Bottom panel illustrate schematic drawings of the fusion events. Scale bar = 5 μm. D. FOXR1 was sequentially extracted with Tris-HCl, Triton X-100, Sarkosyl and SDS in this order. Quantification shows that the amount of FOXR1 in the SDS fraction was significantly higher in the M280L mutant when compared to the overall Tris-HCl total fraction. One-way ANOVA (n = 2 independent experiments, ** p<0.005, *** p<0.0001).

Misfolded proteins often expose their hydrophobic domains, leading to aggregation (Kim *et al*, 2013; Diaz-Villanueva *et al*, 2015). In addition, most aggregated proteins tend to coalesce and form large deposits such as aggresomes or inclusion bodies (Kopito, 2000; Markossian & Kurganov, 2004). Previous studies have shown that nuclear and cytoplasmic aggregates of poly-Q proteins such as ataxin-1 are dynamic and exchange their components whereas ataxin-3 are immobile (Stenoien *et al*, 2002; Kim *et al*, 2016). In fact, time-lapse live cell imaging of HEK293T cells transfected with GFP-tagged M280L showed that the nuclear aggregates are quite dynamic and undergo extensive movements and fusions, with small aggregates moving toward each other and fusing to form larger aggregates (Fig 7C, Appendix movie 3).

Another criterion of misfolded proteins deposited within aggresomes is that they are largely detergent insoluble (Jensen *et al*, 1995; Ward *et al*, 1995; Scherzinger *et al*, 1997; Garcia-Mata *et al*, 1999; Kopito, 2000). Thus, we examined the biochemical properties of M280L aggregates versus FOXR1 WT, testing protein lysates from HEK293T cells transfected with GFP, GFP-tagged FOXR1 WT or M280L for their solubility in different detergents. Protein extracts were sequentially extracted by Tris-HCl buffer followed by Tris-HCl buffer containing 1% Triton-X100, then by 1% Sarkosyl and finally by 2% SDS. The amount of FOXR1 extracted in each fraction was assessed by immunoblotting for GFP-FOXR1. GFP-FOXR1 WT was detected in Tris-HCl soluble, sarkosyl soluble and SDS soluble fractions but not present in Triton X-100, suggesting that the majority of the FOXR1 WT protein was soluble, not associated with membrane-bound proteins (Fig 7D). However, the majority of M280L was detected in the SDS fraction indicating a significant portion of the protein was insoluble and aggregating, which correlates with the increased aggregation shown by the proteostat immunolabelling.

## Discussion

The UDN has identified an individual presenting with severe neurological symptoms and linked a missense variant in the *FOXR1* gene as a potential variant underlying the genetic etiology of the rare neurodevelopmental disorder (Fig 1). The single *de novo* missense variant in *FOXR1* converts a highly conserved methionine residue at amino acid 280 to leucine and was predicted to be damaging and disease-causing based on several web-based applications. Indeed, we found the M280L mutation in FOXR1 leads to a robust decrease in FOXR1 protein expression that is due to protein instability (Fig 2). Protein levels of both FOXR1 WT and M280L were approximately the same after proteasome inhibition, suggesting that the M280L mutant destabilizes FOXR1 protein, likely due to protein misfolding making it susceptible to proteolysis and degradation through the proteasome pathway. To support this finding, we found the M280L variant formed discrete nuclear puncta that colocalize with the Proteostat dye which recognizes misfolded and aggregated proteins compared to the diffuse nuclear pattern localization in FOXR1 WT (Fig 3 and 7). The M280L nuclear puncta displayed characteristic features of most aggregated proteins wherein smaller foci coalesce to form larger aggregates that were biochemically detergent-insoluble impacting on FOXR1 function (Kopito, 2000). We found the C-terminal sequences of FOXR1 are important for determining protein stability. A FOXR1 C-terminal truncation mutant lacking the last 12 amino acids from M280 (Δ280-292) mimics the M280L phenotype, thus suggesting that the M280L mutation most likely affected a conserved amino acid critical for protein stability.

Here we identified target genes regulated by FOXR1 based on an unbiased transcriptomic screen using RNAseq in HEK293T cells (Fig 4). Differential gene expression analysis identified FOXR1 acts as both a transcriptional activator and repressor. The most highly upregulated genes in response to FOXR1 WT and downregulated in M280L included two members of the Hsp70 family (*HSPA1A, HSPA6)* and a mitochondrial reductase enzyme, *DHRS2*. Each of these proteins play a role in protecting against oxidative stress-mediated cellular response. In addition, the top FOXR1-regulated genes *HSPA6, HSPA1A* and *DHRS2* contain FOXR1 response elements within their promoter regions and are direct targets of FOXR1; M280L abolishes its ability to activate the expression of these target genes.

Cells respond to environmental stressors though the activation of specific physiological pathways that increase the abundance or activity of chaperone proteins which prevent protein misfolding to protect the proteome and maintain proteostasis (Voellmy, 1994; Grune *et al*, 2004; Scandalios, 2005; Raynes *et al*, 2016). One important mechanism is the induction of expression of Hsps such as the large Hsp70 family of proteins that help to maintain proteostasis by acting as molecular chaperones during periods of acute cellular stress (Jaattela *et al*, 1992; Gabai *et al*, 1997; Iordanskiy *et al*, 2004). It is well-established that HSF regulates the expression of the Hsps during times of stress where HSF binds to heat shock elements within the promoter regions of Hsps (Nollen & Morimoto, 2002). However, there is now growing evidence that Fox family of transcription factors also influences Hsp expression. For example, the FOXO subfamily of transcription factors play an important role in protecting organisms against stress (Kops *et al*, 2002; Bakker *et al*, 2007; Eijkelenboom *et al*, 2013). Both *FOXO* genes in *Drosophila* (*dFOXO*) and in *C. elegans* (*DAF16*) are transcriptional activators for Hsp70 genes that contribute to maintaining proteostasis in response to oxidative stress. DAF16 maintains proteostasis in *C. elegans* by transcriptionally increasing a subset of small Hsp genes that is important in DAF-16 dependent lifespan extension (Hsu *et al*., 2003; Murphy *et al*, 2003). Consistent with *C. elegans, Drosophila* dFOXO also induces transcription of Hsp genes in response to oxidative stress resistance (Donovan & Marr, 2016). Mammalian FOXO3 and FOXM1 orchestrate programs of gene expression that regulate oxidative stress resistance by upregulating catalase and MnSOD, enzymes involved in the detoxification of reactive oxygen species (Kops *et al*., 2002; Bakker *et al*., 2007; Park *et al*, 2009; Gurkar *et al*, 2018). Here, we show that FOXR1 protein expression is increased in response to metabolic and oxidative stress that increase HSPA6 and DHRS2 protein levels. We demonstrated that HSF1 bind to the FOXR1 promoter and induces its transcription, indicating FOXR1 is a direct target of HSF1. Perhaps HSF is a master transcription factor responding to stress and cross-talk with FOXR1 serves to fine tune transcription of target genes in response to specific stress stimuli.

An outstanding question is what role *FOXR1* plays in the process of brain development. Although a preliminary diagnosis implicating the *ATP1A3* variant for this patient has been made, a synergistic contribution from additional mutations including the *FOXR1* M280L variant cannot be ruled out. Human genetic analyses show that several FOX transcription factors have important biological functions in brain development and mutations in *FOX* genes have profound effects on development and function of the brain. FOXG1, formerly named Brain Factor-1 (BF-1), is one of the earliest transcription factors expressed in the nervous cell types and tissues. FOXG1 is primarily expressed in the telencephalon and *foxg1* knockout mice showed severe microcephaly with a reduction in size of the cerebral hemispheres (Xuan *et al*, 1995). Mechanistically, FOXG1 interacts with the global transcriptional corepressors of the Groucho/transducing-like Enhancer of the split (TLE) family suggesting that FOXG1 acts as a transcriptional repressor coordinating the control of neural progenitor cell proliferation with the timing of differentiation (Yao *et al*, 2000). Disruption in *FOXG1* in humans leads to structural brain abnormalities including microcephaly and agenesis of the corpus callosum (Shoichet *et al*, 2005; Kortum *et al*., 2011; Hettige & Ernst, 2019). In addition, human mutations in both FOXP1 and FOXP2 lead to severe speech and cognitive impairments (Lai *et al*, 2000; MacDermot *et al*., 2005; Takahashi *et al*, 2009; Horn *et al*, 2010; Nudel & Newbury, 2013; Han *et al*., 2019); where both genes have also been linked to autism spectrum disorders (Takahashi *et al*., 2009; Mukamel *et al*, 2011; Bowers & Konopka, 2012b, a).

The RNAseq analysis in HEK293T cells transfected with FOXR1 WT revealed some of the upregulated genes were involved in ribosome biogenesis such as ribosome biogenesis regulator 1 (*RRS1*) and nervous system development (*MTURN, PDZD8, PTPRZ1, NOTCH2*). Since HEK293T cells originate from neural crest cells, this might explain the expression of several neuron-specific genes. Ribosome biogenesis is a key driver in neurodevelopment and dysregulated ribosomal biogenesis result in neurodevelopmental syndromes that present with microcephaly, autism, intellectual deficits and/or progressive neurodegeneration (Hetman & Slomnicki, 2019). Also, ribosome assembly is an energy-demanding process and alteration of any step in ribosomal biogenesis is highly prone to proteotoxic stress that triggers rapid activation of a specific stress pathway that coordinately upregulates heat shock target genes (Albert *et al*, 2019). It is possible that FOXR1 plays a role in protection against proteotoxic stress during ribosome assembly which is essential in brain development. We propose that FOXR1 is a transcription factor that regulates critical genes necessary during brain development which are involved in balancing growth and protein homeostasis. Therefore, understanding how FOXR1 regulates the transcription of genes and how this influences brain development are important questions to address in future experiments.

## Materials and Methods

### Proband enrollment and consent

The proband was evaluated at the National Institutes of Health Undiagnosed Diseases Program (NIH UDP) and was enrolled in the protocol, approved by the National Human Genome Research Institute Institutional Review Board. The parents of the proband provided written informed consent for medical and genetic studies designed to reach medical diagnoses.

### Exome sequencing

Exome sequencing was performed using genomic DNA extracted from peripheral whole blood samples from the study participant and family members after informed consent onto an institutional review board approved protocol (76-HG-0238). Exome capture was carried out using manufacturer protocols using the TruSeq Exome Enrichment Kit (Illumina, San Diego, CA) and sequenced on the HiSeq 2000 Sequencing System (Illumina). Alignment to the human genome reference sequence (UCSC assembly hg19, NCBI build 37) was carried out using the Efficient Local Alignment of Nucleotide Data algorithm (*Eland*, Illumina, Inc) as described previously (Yuan *et al*, 2014). Briefly, paired-end (PE) reads were aligned independently and reads that aligned uniquely were grouped into genomic sequence intervals of ∼100 kb whereas reads that failed to align were binned with PE mates without *Eland* using the PE information. Reads that mapped in more than one location were discarded. To align binned reads to their respective 100 kb genomic sequence, *Crossmatch*, a Smith-Waterman-based local alignment algorithm was used based on the following parameters –minscore 21 and –masklevel 0 (http://www.phrap.org). Genotypes were identified using a Bayesian genotype caller, Most Probable Genotype (Teer *et al*, 2010). Selected *de novo* variants detected exclusively by exome sequencing were tested by Sanger sequencing. Sanger sequencing was performed to confirm the segregation of the identified variant in *FOXR1* using the following primers F, 5’-AAAGCACTTCCCCTTTTTCC-3’ and R, 5’ AGTTGTTTGCCCATGGATTC-3’.

### Construction of expression vectors

Full-length human pCMV-SPORT6 *FOXR1* plasmid was purchased from GE Dharmacon (clone ID 5164198; accession #BC038969). The human M280L mutation in *FOXR1* was generated by introducing a point mutation at residue 280 (methionine to leucine) using QuikChange II Site-Directed Mutagenesis Kit (Agilent Technologies) in the pSport6 human FOXR1 plasmid with the following forward 5’-CCAACAGTGCTTGAGCCAGCCAG-3’ and reverse 5’-ATACTTTCTAGCCGAGTGGAAG-3’ primers and verified by nucleotide sequencing. The *FOXR1* wild-type and M280L mutant were then PCR amplified using forward 5′-AAAGCACTCGAGATGGGGAACGAGCTCTTTCTG-3’and reverse5’-TTTGGCCCGCGGTTAAAGATCAAAGAGGAAGGG-3’ primers and subcloned into the XhoI and SacII restriction sites of pEGFP-C3 (Clontech) to create an N-terminal EGFP tag. To generate the *FOXR1* C-terminal truncation variant, Δ280-292, we used full-length human GFP-tagged *FOXR1* wild-type as template and designed PCR forward 5’-AAAGCACTCGAGATGGGGAACGAGC-3’and reverse5’-TTTGGCCCGCGGTTAGCACTGTTGGATACTTTCTAGCCG-3’ primers to amplified amino acids 1-279 and subcloned into the XhoI and SacII restriction sites of pEGFP-C3.

### Cell culture

HEK293T (ATCC CRL-3216) and COS-7 (ATCC CRL-1651) cells were maintained in Dulbecco’s Modified Eagle’s Medium (DMEM) supplemented with 10% fetal bovine serum (FBS; Hyclone) and 1% Penicillin/Streptomycin at 37°C in a 5% CO_2_ incubator. Cells at 60% confluency were transfected with GFP, GFP-FOXR1 or GFP-M280L plasmids using FuGENE6 transfection reagent (Promega) according to manufacturer’s instructions. For subcellular fractionation, cells were briefly washed with phosphate-buffered saline (PBS) and lysed in buffer A that consists of 50 mM Tris-HCL pH 7.5, 0.5% Triton-X100, 137.5 mM NaCl, 10% glycerol, 5 mM EDTA pH 8.0 with proteinase inhibitors. The lysate was centrifuged 850 x *g* for 15 min at 4°C. The supernatant “cytosolic” fraction was removed to a new tube and the remaining “nuclear” pellet was washed twice with buffer A at 4°C and centrifuged at 850 x *g* for 2 min. The pellet was then solubilized in buffer B that consists of 50 mM Tris-HCL pH 7.5, 0.5% Triton-X100, 137.5 mM NaCl, 10% glycerol, 5 mM EDTA pH 8.0, 0.5% SDS with proteinase inhibitors and sonicated for 5-10 secs. Equal amount of 2x sample buffer (0.1 M Tris-HCL, pH 6.8, 4% SDS, 20% glycerol, 10% β-mercaptoethanol, 0.01% bromophenol blue) was added to the tubes containing the nuclear and cytoplasmic fractions, boiled at 100°C for 10 min and subjected to SDS-PAGE. For MG132 treatment, transfected cells were treated with 50 μM MG132 (Sigma-Aldrich) for 24 h. Cells were then washed with PBS and collected in 2x sample buffer. Cellular stress paradigms: serum starvation, cells were incubated in DMEM without fetal bovine serum for 24 h at 37°C; for CO_2_ deprivation, cells were deprived of 5% CO_2_ for 24 h at 37°C; PMA treatment, cells were treated with 1 μM of phorbol 12-myristate 13-acetate (PMA, Sigma) for 24 h at 37°C.

### Western blotting

Whole cell lysates were extracted from cells in 2x sample buffer and separated on 10% SDS– PAGE gel and transferred to nitrocellulose membranes (GE Healthcare). The membranes were blocked with Odyssey Blocking Buffer in PBS (Licor), followed by incubation with primary antibodies against human FOXR1 (Biorbyt, 1:200), GFP (synaptic systems, 1:1000), GAPDH (EMD Millipore, 1:5000), HSPA1A and HSPA6 (Enzo life sciences, 1:1000), DHRS2 (Abcam, 1:500), Histone H3 (Cell-Signaling, 1:1000) overnight at 4°C. Proteins recognized by the antibodies were detected with an Odyssey infrared imaging system (LI-COR) using IRDye680RD- or IRDye800CW-coupled secondary antibodies (LI-COR, 1: 20,000).

### Immunocytochemistry and image analysis

Transfected cells plated on coverslips were washed briefly with PBS and fixed with 4% paraformaldehyde at room temperature for 10 min, permeabilized and blocked in 10% goat serum, 0.1% saponin in PBS. To detect oxidative stress following PMA treatment, transfected cells were incubated with 5 μM CellROX Oxidative Stress Reagent (Thermo Fisher Scientific) for 30 min at 37°C prior to fixation with paraformaldehyde. To detect aggresomes in transfected cells, we used the PROTEOSTAT Aggresome Detection Kit (Enzo Life Sciences). The dye intercalates into the cross-beta spine of quaternary protein structures found in misfolded and aggregated proteins. Coverslips were mounted with ProLong Gold Anti-Fade Mount with DAPI (Fisher Scientific) and imaged with a Carl Zeiss LSM700 confocal microscope. Images were collected with identical confocal settings for all of the samples and Z-stacked images were projected with maximal projection mode using Zeiss Confocal Software.

### RNA sequencing and analysis

HEK293T were transfected with GFP, GFP-FOXR1 or GFP-M280L mutant using FuGENE6. Forty-eight hours after transfection, total RNA was purified using the QIAshredder and RNeasy Mini Kit (QIAGEN) and samples were processed with Trizol (Invitrogen). Three biological replicates were processed independently. RNA samples were suspended in DEPC-treated water and concentrations were determined using the Nanodrop ND-1000 (Thermo Scientific) where all samples showed A260/A280 ratios higher than 2.0 and RNA integrity were also checked in a bioanalyzer (Agilent 2100). Library preparations and sequencing were performed by The Broad Institute, MA using Illumina HiSeq 2000 technology.

The RNA sequencing reads were aligned to the GRCh38 *Homo sapiens* genome using HISAT2 (Kim *et al*, 2019) with default parameters. The bam files were sorted by read names instead of chromosome coordinates by SAMtools (Li *et al*, 2009). Gene count matrix of each sample was generated by HTSeq, a Python framework to work with high-throughput sequencing data (Anders *et al*, 2015). Downstream analysis was performed with the DEseq2 (Love *et al*, 2014) package in R. Gene that were not expressed in any cell and were removed from downstream analysis. Sample PCA plot was generated with ‘plotPCA’ function to detect and remove the outlier sample in each condition. Differential expression analysis between conditions was performed with the ‘DESeq’ function with default parameters. Log-fold change shrinkage was performed on the differential expression analysis result. DEGs with adjust-p value < 0.05 and log-fold-change > 0.25 were kept for downstream analysis. Heatmap of differentially expressed genes were visualized with heatmap and genes with similar expression patterns across samples were clustered on the heatmap. Gene set enrichment analysis was conducted with the GSEA (Subramanian *et al*, 2005). Gene ontology and enrichment analysis encompassing the DEGs were analyzed using the Database for Annotation, Visualization, and Integrated Discovery (DAVID v6.8) software where the threshold was set as modified Fisher Exact *P*-value (EASE score) ≤ 0.05.

### Quantitative real-time qPCR analyses

Total RNA was purified using the QIAshredder and RNeasy Mini Kit (Qiagen). The cDNA was synthesized using iScript cDNA Synthesis Kit (Biorad) or Accuris qMax cDNA Synthesis Kit (Midland Scientific). Quantitative real-time PCR was performed in an ABI Prism 7900HT Fast Real-Time PCR System (Applied Biosystems) using the Power SYBR Green PCR Master mix (Thermo Scientific) with a two-step cycling protocol and an annealing/extension temperature of 60°C. The experiment was performed with three biological replicates and three technical replicates each. The relative amount for each target was normalized using GAPDH as a reference gene and the fold change in gene expression was calculated using the ddCt method with the GFP-transfected cells serving as control. Primers were as follows: *DHRS2*: 5′-TCATCAGCTGCAGAGGATTGG-3′ (forward) and 5′-AATGTTCTCCCCGTTGACGTA-3′ (reverse); *DNAJC6*: 5’-AGGACAACTTGAAAGACACCCT-3’ (forward) and 5’-AAATCTCCCTTTGTGTAGCTGG-3’ (reverse); *DNAJC21*: 5’-CCTGAAATGGCACCCGGATAA-3’ (forward) and 5’-TTTCCTGAGGGTCACTCAACA-3’ (reverse); *GAPDH*: 5′-GGATTTGGTCGTATTGGG-3′ (forward) and 5′-GGAAGATGGTGATGGGATT-3′ (reverse); *HSPA1A*: 5′-GCCTTTCCAAGATTGCTGGTT-3′ (forward) and 5′-TCAACATTGCAAACACAGGA-3′ (reverse); *HSPA6*: 5′-CAAGGTGCGCGTATGCTAC-3′ (forward) and 5′-GCTCATTGATGATCCGCAACAC-3′(reverse);*HSPA12A*:5’-GCTCCCACATCTGCATATTCAT-3’ (forward) and 5’-TTCTGAGACGTTGGAGTCAGT-3’ (reverse); *SACS*: 5’-ACAACAACGCGGTTTTCACC-3’ (forward) and 5’-GCCTGATTCATGTGGGCCAA-3’ (reverse). Data analysis was performed using the ABI Prism 7900HT SDS Software.

### Dual luciferase assay

Promoter sequences of human *HSPA1A, HSPA6, DHRS2* were amplified by PCR from genomic DNA and subcloned into pNL3.1-minP/Nluc (Promega). Cells were transfected with the normalization plasmid (pGL4.54-Luc2/TK), the reporter plasmid (pNL3.1-minP/Nluc) and GFP-tagged FOXR1 or M280L plasmid. Transfected cells were collected in PBS and luciferase activity was assessed using the Nano-Glo Dual Luciferase reporter assay system (Promega). Dual luciferase signal was quantified using a VICTOR-3 plate reader (Perkin Elmer). To control for transfection efficiency, the Nluc reporter plasmid signal was normalized to the constitutive luciferase signal (i.e., signal from pGL4.54 plasmids, Nluc/Luc2). Fold-induction values for each promoter sequence were calculated relative to the background activity of reporter plasmid in the presence of GFP-FOXR1 or GFP-M280L plasmid. Reporter assays were performed as three biological replicates with three technical replicates per biological replicate.

### Detergent extraction assay

Transfected HEK293T cells were isolated in 50 mM Tris-HCl buffer at pH 7.5 with protease inhibitors by brief sonication and ultracentrifuged at 350,000 x *g* for 15 min and the supernatant was collected as a Tris-HCl soluble fraction (adapted from Kuwahara *et al*, 2012). The resulting pellet was sequentially extracted in Tris-HCl buffer containing 1% Triton X-100, then by 1% Sarkosyl and finally by 2% SDS. Each detergent extraction step was incubated for 1 h at 4°C and ultracentrifgued at 350,000 x *g* for 15 min, resulting in a Triton X-100 soluble fraction, Sarkosyl soluble fraction and SDS soluble fraction, respectively. The Tris-HCl fraction containing 20 < g of total proteins, along with equal volumes of Triton X-100, Sarkosyl and SDS fractions were loaded onto SDS-PAGE.

### Statistical analysis

To determine statistical significance, we use either a Student’s *t*-test to compare two groups, or a one-way repeated measures ANOVA for multiple comparisons and two-way ANOVA for comparisons with multiple variables. All bars and error bars represent the mean <u>±</u> S.E.M. and significance was set at p<0.05. The data were analyzed using GraphPad Prism 7 software (GraphPad Software Inc., San Diego, CA).

## Data availability

The accession number for the RNAseq reported in this paper is GEO:GSE

### Acknowledgements

We thank the colleagues in the laboratory who have provided constructive feedback on this work. This work was supported in part by the Intramural Research Program of the National Human Genome Research Institute (HG000215 to W.A.G.) and by a National Institutes of Health grant (R21GM114629 to U.B. and A.H.).

## Author Contributions

A.M., A.H., U.B. Conceptualization; A.M., R.H., S.N., C.C., L.W. Data curation; A.M., R.H., T.M. Formal analysis; A.H., U.B., W.A.G. Funding acquisition; A.M., R.H., S.N., C.M.M., C.C, U.B., A.H. Investigation; A.M., R.H., S.N., F.B., D.R.A., Methodology; D.R.A., W.A.G., C.C., U.B., A.H. Project Administration; C.M.M., D.R.A., W.A.G., C.C., U.B., A.H. Supervision; A.M., R.H., T.M., Validation; A.H. Writing-original draft; A.M., C.M.M., W.A.G., U.B., A.H. Writing-review & editing

## Appendix figure legends

**Appendix Table 1**. Summary of clinical features of the proband with a *de novo* variant in *FOXR1*.

**Appendix Figure 1**. Alignment of cross-species FOXR1 sequences showing the conserved methionine residue (indicated in red) within a highly conserved region of amino acid sequence.

**Appendix Figure 2**. Fluorescence images of COS7 cells transfected with GFP or GFP-tagged plasmids of human FOXR1 WT or M280L. DAPI (blue) served as a nuclear marker. Scale bar = 20 μm.

**Appendix Figure 3**. RNAseq analysis

A. PCA plot of the three groups clustered separately in multidimensional scaling analyses. Groups of samples analyzed using Principal Component Analysis (PCA) plots where replicates are clustered together and clusters from different conditions are separated.

B. B, Pie chart showing the distribution of 2644 differentially-expressed genes between GFP and FOXR1 WT and 735 differentially-expressed genes between FOXR1 WT and M280L.

**Appendix Figure 4**. Gene ontology (GO) enrichment analysis between FOXR1 WT and M280L and GFP control and FOXR1 WT. Normalized enrichment scores indicate the distribution of biological processes across a list of genes ranked by hypergeometrical score. Higher enrichment scores indicate a shift of genes belonging to certain GO categories towards either end of the ranked list, representing up or down-regulation (positive or negative values, respectively).

**Appendix Figure 5**. Promoter sequences showing *FOXR1* response elements lie upstream of the HSEs in both the *HSPA6* and *HSPA1A* promoters. In addition, we identified a consensus sequence for binding by HSF1 within the promoter region of *FOXR1*.

**Appendix Movie 1**. Movie of M280L in response to CO_2_ stress.

**Appendix Movie 2**. Movie of M280L in response to PMA treatment.

**Appendix Movie 3**. Movie of M280L in response to PMA treatment at high magnification.

