## Appendix Fig 1 for "Forkhead box R1-mediated stress response linked to a case of human microcephaly and brain atrophy"

| | | | |
|---|---|---|---|
| Human | 249 | CLWKLTEEGHRRFSKEARTLASTQLQSIQQCMSQPGVKPFLFDL | 292 |
| Chimpanzee | 249 | CLWKLTEEGHRRFAEEARALASTRLESIQQCMSQPDVMPFLFDL | 292 |
| Rat | 227 | CLWKLTEEGHRRFSEEARTLASTRLQSIQQCMSQPGVKP----- | 265 |
| Mouse | 225 | CLWKLTEEGHRRFSKEARTLASTQLQSIQQCMSQPGVKPFLFDL | 268 |
| Bovine | 212 | CLWKLTDEGHRRFAEEARALASTRLERIQQCMSQPDVMSSLFDL | 255 |
| Goat | 248 | CLWKLTDEGHRRFAEEARALASTRLERIRQCMSQPDVMSSLFDL | 291 |
| Horse | 250 | CLWKLTEEGHRRFAEEARALASTRLESIQRCMSQPDVMPFLFDL | 293 |
| Panda | 281 | CLWKLTKEGHRRFAEEARALASTHLESIQQCMSQPDVMPFLFDL | 324 |
| Walrus | 254 | CLWKLTEEGHRRFAEEARTLASTRLESIQQCMSQPDVMPFLFDL | 297 |
| Dolphin | 249 | CLWKLTDEGHRRFAEEARALASTKLETIQQCMSQPDVMPSLFDL | 292 |
| Fox | 253 | CLWKLTEEGHRRFTEEARALASTRLQSIQQCMSQPDVMPFLFDL | 296 |
| Dog | 251 | CLWKLTEEGHRRFVEEVRALASTRLESIQQCMSQPDVMPFLFDL | 294 |
| Cat | 252 | CLWKLTEEGRRRFAEEARALASSRLESIQRCMSQPDVMPFLFDL | 295 |
| Pig | 249 | CLWKLTEEGHRRFVEEVRALASTRLESIQQCMSQPDVMPFLFDL | 292 |
| Rabbit | 269 | CLWKLTEEGRRRFAEEARALAATQLGSIQQCMSQPDMMPFLFDL | 312 |
| Ferret | 280 | CLWKLTEEGHRRFAEEARALVSTRLESIQQCMSQPDVMPFLFDL | 323 |
| Armadillo | 273 | CLWRLTEEGHRRFQEETRALASARRESIQQCMSRPDVMTFLFDL | 316 |
| Bat | 268 | CLWKLTEEGRRRFEEEARALSSAQLERIQQCMSQPGVMPFLFDL | 311 |
| Penguin | 261 | CLWKLTPEGRRKFQEEAQALPKEALDLVRQSMSEPELMRSLFGL | 304 |
| Alligator | 255 | RLWKLTTDGRRKFEEEMQALPTEALALVHQSMNEPELMSSLFCL | 298 |
| Python | 110 | RLWKLTTEGRRKFQEEAQALPEDMMGLVHQSMDKPELMRSLFGL | 153 |
| Frog | 196 | CLWKLTRQGRRKFRNEMHALSDDLLHVLRKSMKKPALMELMFGM | 239 |
| Zebrafish | 278 | CLWHLTLDGRQRLRDEIHTLTEDSFRLLRRSMNYPDMIPALLEL | 321 |
