## Supplementary figures and images for "Forkhead box R1-mediated stress response linked to a case of human microcephaly and brain atrophy"

### Appendix Fig 2

#### Appendix Fig 2

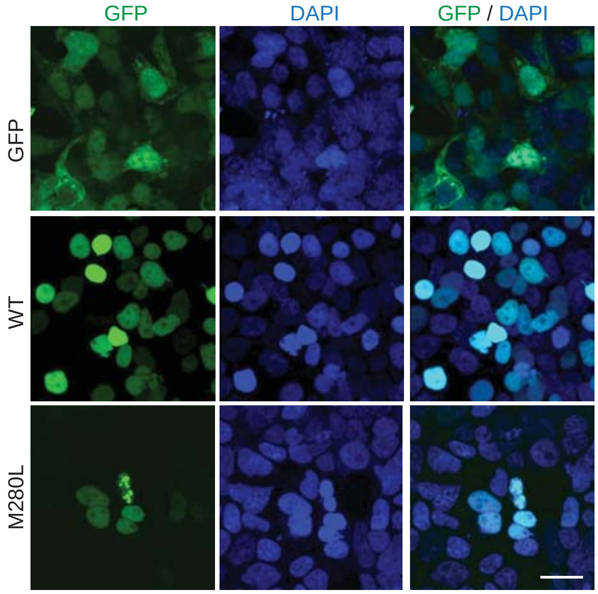

### Appendix Fig 3

#### Appendix Fig 3

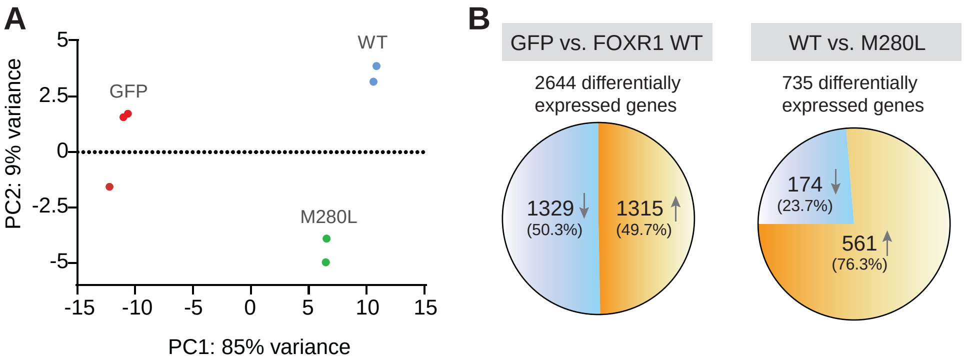

### Appendix Fig 4

#### Appendix Fig 4

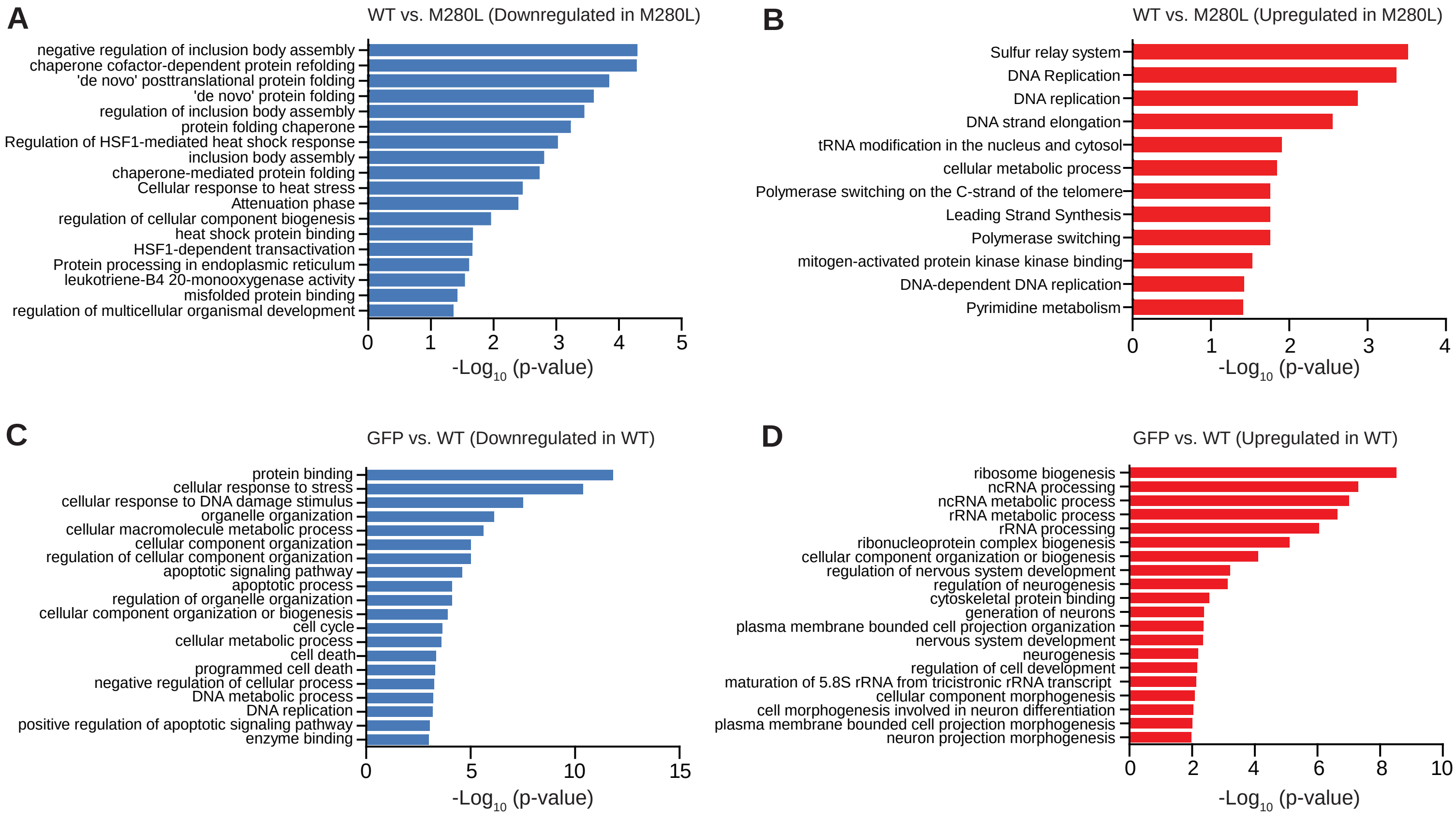
