## Appendix Fig 5 for "Forkhead box R1-mediated stress response linked to a case of human microcephaly and brain atrophy"

FOXR1 Binding site (Primary CAAACA/ Partial GACGCC)

HSEs

NNNNN represents nucleotides not cloned into luciferase plasmid

TATAAA represents TATA box

**Human *HSPA1A* Promoter (-1043 →-152)**

TCCGACCAATCAATCTGAAGCCATCTTAGCTTTCCCCAAGTGCTCCTCCTACCCGGATCAG
CCAACGCCCACATACCTCAGGCTTAAACCAACTAGGGAACTTTCCAGTACTTTCCCAAACA
AGGACCTACTGAGCCTTTCAGGTTCACAATCAATCAGATCCCTACTGGCTCACCTAGTCTC
CCGACGCCTTCGCTTCAGTTTGGAAACGTCCAGATTACGCAGCCCCAGCGAGTAGGTGGG
GGCTCCCTCAATATCAAACTGCACAACCGGGGTCCCCCCACCCCCCACCCCGTCCCTCCC
TGCAAATTTGAGACGGCTCCAACTCAGTAATCTTTTTCCAAACTGGCCCATGAGGTCAGAG
ACAGTATCTCCATTGTAACGTGGCCGGGCGGTGTCAACACAAACGCCCCCACCCTCCCCT
GGACGCGCGTAACCCGCTCCCCGCACCAGCCCCCTGCCCACAACTGCGCAGGCCCAGCA
AGCCCCCACAATTAAAAGCCCAGCGCCGACCCTTCCTGTCAATTAGGCGCTGAAGCGCAG
GCGGTCAGCATCGCCATGGAGACCAACACCCTTCCCACCGCCACTCCCCCTTCCTCTCAG
GGTCCCTGTCCCCTCCAGTGAATCCCAGAAGACTCTGGAGAGTTCTGAGCAGGGGGCGG
CACTCTGGCCTCTGATTGGTCCAAGGAAGGCTGGGGGGCAGGACGGGAGGCGAAAACCC
TGGAATATTCCCGACCTGGCAGCCTCATCGAGCTCGGTGATTGGCTCAGAAGGGAAAAGG
CGGGTCTCCGTGACGACT**TATAAA**AGCCCAGGGGCAAGCGGTCCGGATAACGGCTAGCC
TGAGGAGCTGCTGCGACAGTCCACTACCTTTTTCGAGAGTGACTCCCGTTGTCCCAAGGC
TTCCCAGAGCGAACCTGTGCGGCTGCAGGCACCGGCGCGTCGAGTTTCCGGCGTCCGGA
AGGACCGAGCTCTTCTCGCGGATCCAGTGTTCCGTTTCCAGCCCCCAATCTCAGAGCGGA
GCCGACAGAGAGCAGGGAACCGGC**ATG**

**Human *HSPA6* Promoter (-1119→-114)**

CACCGGGCCTCTGGAGACGAGGCTCCTCGGGGATACAAACAGTGGGGAGAACATGAGGG
ACATCCCGACCGTACTCTGCGTCCTCCTTTCCCAGGTGTTGCGTTCTCTCTTGGGCTGAGT
GGCGAGGTCTCTCCCGAGTCCCAGGGCCACAGTGCAATGTCACATCTCCTTTGTGGAAAG
TGACTGGTAAAGGAGAGAGAACAAAACTGGAGGAATGTAAAGTCTTCAGCCACCTGGTTTA
ATTTATTCAAGAGTGATTAATCCTAGATGAGAAAAAGAATTGAAATGGATCGGAAAAAAATG
AAAGTGCATTGGCCGGGAATCGAACCCGGGCCTCCCGCGTGGCAGGCGAGAATTCTACC
ACTGAACCACCAATGCTACTGTCAGCTAAAGACCTGCAGTATTGTCTCTTAAAGCTCACTAT
CTCTGGCCATTCACTAAGGAACCAGGCACCGTCTTAAATCGCGGTTTGGAAAATATTTTGTT
CAAGATAAAACTGTTTTAAGATATACGTGTATATATCTTATATATCTGTATTCGCATGGTAAC
ATATCTTCGGCCTTCCTGAGCCGCTGGGCTCTCAGCGGCCCTCCAAGGCAGCCCGCAGG
CCCCTGTGTGCCTCAGGGATCCGACCTCCCACAGCCCCGGGGAGACCTTGCCTCTAAAGT
TGCTGCTTTTGCAGCCTCTGCCACAACCGCGCGTCCTCAGAGCCAGCCCGGAGGAGCTAG
AACCTTCCCCGCATTTCTTTCAGCAGCCTGAGTCAGAGGCGGGCTGGCCTGGCGTAGCCG
CCCAGCCTCGCGGCTCATGCCCCGATCTGCCCGAACCTTCTCCCGGGGTCAGCGCCGCG

CCGCGCCACCCGGCTGAGTCAGCCCGGGCGGGCGAGAGGCTCTCAACTGGGCGGGAAG
GTGCGGGAAGGTGCGGAAAGGTTCGCGAAAGTTCGCGGCGGCGGGGTCGGGTGAGGC
GCAAAAGG**ATAAAAA**GCCGGTGGAAGCGGAGCTGAGCAGATCCGAGCCGGGCTGGCTG
CAGAGAAACCGCAGGGAGAGCCTCACTGCTGAGCGCCCCTCGACGGCGGAGCGGCAGC
AGCCTCCGTGGCCTCCAGCATCCGACAAGAAGCTTCAGCC**ATG**

**Human *DHRS2* promoter sequence (Partial) (-3329 → -2313)**

TAATGCCAAATCATTTCCCAAAGTGATTGTACTTACCCTCTACCCAGCAGTGAATGAGAGTT
TCTGTTGATTTTGCTTTTTCAACTAGAAGTGAGAGGTGACAATATGCTAGCAGCCCTTGCTC
ACTCTTGGTGCCTCCTCGTCCTCCACATCCACTCTGGCTGCACTTGAGGAGCCCTTCAGCC
CACCACTGCACTGTGGGGGCCCCTCTCTGGGGCTGGCCAAGGCCGAAGCCAGCTCCCTC
TGCTCGCCAGGAGGTGTGGAGGGAGAGGCGCCGGCAGGAGCCCACACTGTACAGGGCA
CTCACCGGCCAGCAGGGGCTCCGTGGGCCGGCCGGTGCCAGCTGGGCCTGACTGGGGG
ATGAGCTCCCTCTGGGCTGCTGGAGTGCCTGGGCTAGGTGCCGCAAAGTCCTGCAGTGT
GTGCCATTGAGAGGTGAAGCCGGCTGGGCTTCTGGGTCCGGTGGGGACCTGGAGAACTT
TTCTGTCTAGCTAAAGGTTTGTAAATGCACCCATCAGCACTCTGTGTCTAGCTAAAGGTTTG
TAAATGCACCAATCAGCAATCTGTGTCTAGCCAATCTGGTGGGGACTTGGAGAACTTTTGT
GTCTAGCTAAAGGATTGTAAATGCACCAATCAGCACTCTGTGTCTAGCTAAAGGTTTGTAAA
TGCACCAATCAGCACTCTGTCAAAACGGACCAATCAGCTCTCTGTAAAACAGACCAATCAG
CTCTCTGTAAAATGGACCAATCAGCTCTCCATAAAATGGACCAATCAGCAGGATGTGGGTG
TGGCCAGTTAAGGGAATAAAAGCAGGCTGCCTGAGCCGGCAGCAGCAACCTGCTCTGGTT
CCCTTCCACGCTGTGGAAGCTTTGTTCTTTTGGTCTTCATGATAAATCTTGCTGCTGCTCAC
TCGTTGGGTCCGTGCCACCTTTAAGAGCTGTAACACTCACCGCGAAGGTCTGCAACTTCAC
TCCTGGGGCCAGCAAGACCACGAATGCACCGAGAGGAATGAACAACTCTGGACACACCAT
CTTTAAGAACCGTAATACTCACCGCAAGGGTCTGCAACTTCATTCTTGAAGTCAGTGAGGC
CAAGAACCCATCAATTCCGTACACATTTTGGTGACTTTGAAGAGACTGTCACCTATCACCAA
GTGGTGAGACTATTGCCAAGCAGTGAGACTATTGCCAAGTGGTGAGACCATCACCAAGCG
GTGAGACTATCACCTATCGCCAAGTGGTGAGTACCATCAGATCCCTTTCATTTGCTATTCTG
TCCTATTTTTCCTTAGAATTCGGTGGCTAAATTCTGGGCACCTGTCGGCCAGTTAAAAGTGA
CTAGCGCAGCCGCTGGACTAAAAACGTGGGTGTCAGGCTTTCTGGGAAAGGGATCTCTAA
CAACCCCTGGCTCTGTGGAGTTGGGAATGTTGTTTTGCCTGGAACCAGCTTCCGCTTTTCC
TGTACTTCTGGGCTGAGCTGAGGGTCAACAGAGAGGAAAGCCATTCAGCTCCGGAGTCCC
CACAACAAGTTGGTTGACCCTGCGGCCATGAGCGGAACTCTCAAAGGCATGTTGCCCAAG
TGAGACTCACCTATCTATCCTATCTATCCTGACCCTTGCTCGCTGGGTCCTAATGCCTGCC
AGACAAACTTTCTCTCTCCTCTCTTCTCCTAGGCTAGTCCCACTTCTAAAAAACCACTCCCT
GTCTCTGGTGCTTTTCTAATTTCTCTTATAAGAATGATTTCTAGTAAAAATTTCAGGACTCTG
TTACCTTCTTTAGGCACCTGGGCTCACCAATCAGAAAGACATAATTTTTGCCCAAAGCCCA
GTTGTAGGGGGAACTATCTGGAATTTTAGAATCCCTCCTCAGATAAGCAGGTCTAACAAAA
GCTATTCCTGAAGCTAGGATAGGGGGAGCCTCAGAAATTGTATCCTTCCTATTCATATAAGT
GAGGACAAAAGGTGTCACTTTTCCAACCCTGGAGACCCCTTCCCTCCCTCAGGTTACTCTT
CTTCATTTTTGGGGCATAACATCTTTATAGGACACGGGTAAGTTCCCAATACTAACAGGAGA
ATGCTTAGGACTCTAACAGGTTTTCGAGAATGCGTCGGTAAGGGCCACTAAATCCGATTTT
TCTCGGTCCTCCTCATGGTCTAGGAGGACAAGCAAGGGTGCAGCACTCTATGTCTAGCTAA
TCAGGTGGGGACTTGGAGAACTTTTGTGTCTAGCGAAAGGATTGTAAATGCACCAATCAGC
ACTCTGTGTCTAGCTAAAGGTTTGTAAACACAGCAATCAGCACTCTGTCAAAATGGACCAAT

CAGCTCTC**TATAAA**ACAGACCAATCAGCTCTCTGTAAAATGGACCAATCAGCAGGATGTGG
ACAGGGCCAGATAAGGGAATAAAAGCAGGCCACCCGAGCCAGCAGCAGCAACCTGCTTG
AGTCCCCTTCCACACTGTGGAAGTTTTGTTCTTTCGCTCTTTGCAATAAATCTTGATTCACT
GCTGCTCACTCTTTGGGTCCGCGCTGCCTTTAAGAGCTGTAACACTCACCACGAAGGTCTG
CAACTTCACTCCTGAGGCCAGTGAGACCACAAACCCTCCAGAAGGAATGAACAACTCCAG
ACGCACTGCCTTTAAGAGCTGTAACACTCACTGCGAAGGTCTGCAGCTTCACTCCTGAAGC
CAGCAAGACCACTAACCCACCAGGAGGAATGAACAACTCCGGACGGGAGGAATGAACAAC
TCCGGATGGGAGGAATGAACAACTCCGGACACACCATCTTTAAGAACTGTAACACTCATTG
CGAGGGTCCGTGGCTTCATTCTTGAAGTCAGCGAGACCAAGAACCCACCAATTCCAGACA
CAGAAGCAGTTAATTAATAACTGGTATAATTATTTGTTGGTGCCTGGCATCTCATTTAACTG
GGAAGCTGCATGGTTGGGGTCCAGGGCTGAGTCTGTCGTTTACTAAGGTGTCCCCGCCTC
TACTGCCCAGTGCTTGACACATTGCCAGTGCTCAATGTTTGCTATGCAATGGATGGAAAAA
TCACCCTTGTCTAATGAATGTTGAGTCTCACATTTTAATTTGTGAATAATTTCCCCAGTTTTA
TAGCTTAGGAGTTTTCATGGATTGCTTTCCTGACCTGAGGTTCACATGTTTGAAATTTACCC
TAACCAGCCTGACTCTCTGCCACTTTCTGTTGCTGGCCTTGTCTGTCTGGAGGAAGGAGGA
GGGTAGATTACCTTCATGCTCACTGAGGCATCAGTGATAAGTGAAATTGATTCTTTCCCCCA
GGCCTGATTCAGCAGGAAGCATCTCAGACACCAACCACT**ATG**

**FOXR1 promoter region (632bp)**

**HSF1 binding site (-857→ -226)**

GTCCCCCAGGCTGGAGTGCAGTGGCACCATCTCGGCTCACTGCAACCTCCACCTCCCGG
GTTTAAGCGATTCTCCTGCCTCAGCCTCCCTAGTAGCTGGGATTACAGGCGTGAGCCACC
ACGCACTGCTAATTTTTTTGTATTTTTAGTAGAAACGGGGTCTCACCATCGGTCAGGCTGGT
CTCAAACTTCTGACCTCAGGTGATCCGTCCACCTCGGCCTCCCAGAGTGTTGGGATTACAT
GCGTGAACCACCGCATGCGGCCAAAAGACAAAAAAAAATTTTTTTAATCAAATCACCATCCA
ATGTAATTCCATCTCCACCATCTACTACAGATGTGATCGTGGATAAACCATAAACCTTTCAT
AACTTCTAGAAAGTAACTACTAATACTATTATGGTAGAAAGGATCAATGTTAAAGCAAAAC**T**
**ATAAA**AATAAATAAATACGATCACTTGACCATGGCGCCTTTTCAATGAAATAAGGGGCAAG
GGGCTTGGGCAAACGAGGATGCGTAAGATGATCTGGGGTCCCTTCCGGATCCCCAGGTC
CTCGACCCCCGGCATGTAAGGGTAGAGTCAAGGAGGGTCTTCTTAAGCGTTCCTCGGCGC
GGTCCTGAAGAGATTAAAGGCGTCAAATGGACCGCTCCACACCTGAGCTGCCGCCAAGCT
AACTCAATCCGAGCCGGCGCATTTGAGAAGGCGCCTGTGAGGGTCGCTCCTCAGCCGCC
GCGCTCCCACTCCGCGTCCCCACTCCGCGCCGCCGCGCCTCTGCCAGCCCCGAAGGTGG
ACGTGAAGCTCCAACACCTCGACTTCTGGGCCCGCCTCCATGGCCAGGTCCCGGGACTG
CTGGACTGGGAC**ATG**
