## Appendix Table 1 for "Forkhead box R1-mediated stress response linked to a case of human microcephaly and brain atrophy"

| Neuro Phenotypes | Cerebellar Abnormalities | Physical Abnormalities | Facial Abnormalities | Muscle Abnormalities | Visual Abnormalities | Other |
|---|---|---|---|---|---|---|
| Global developmental delays | Cerebellar hypoplasia | Hip dysplasia | Myopathic facies | Hyperactive deep tendon reflexes | Optic atrophy | Exaggerated startle response |
| Postnatal Microcephaly | Abnormality of the cerebellar penduncle | Growth Delay | Preauricular pit | Joint hypermobility | Cortical visual impairment | Aspiration |
| Cerebral white matter atrophy | Cerebellar atrophy | Decreased body weight | Anteverted nares | Severe muscle hypotonia | Retinitis pigmentosa | Constipation |
| Abnormality of midbrain morphology | Aplasia/Hypoplasia of the cerebellum | Short stature | Low set ears | Generalized hypotonia/ dystonia | | |
| Abnormality of the medulla oblongata | | Ankle Clonus | | Muscular hypotonia of the trunk | | |
| Dandy –Walker malformation | | Bell-shaped thorax | | Poor head control | | |
| Hypoplasia of the pons | | Bilateral single transverse palmar creases | | Decreased fetal movement | | |
| Neuronal loss in the cerebral cortex | | Scoliosis | | | | |
| EMG: Neuropathic changes | | | | | | |
| EEG with generalized slow activity | | | | | | |
| Low CSF 5-methyltetrahydro-folate | | | | | | |
| Dilation of lateral ventricle | | | | | | |
| Diffuse cerebral atrophy | | | | | | |
